# *De novo* nitric oxide synthesis drives tactile hypersensitivity induced by ATP-sensitive potassium channel opening in mice: Relevance to migraine and other headache disorders

**DOI:** 10.1101/2025.08.15.670471

**Authors:** Rikke Holm Rasmussen, Charlotte Ernstsen, Anja Holm, Sabrina Prehn Lauritzen, Karina Obelitz-Ryom, David Møbjerg Kristensen, Inger Jansen-Olesen, Jes Olesen, Sarah Louise Christensen

**Affiliations:** Department of Neurology, Danish Headache Center, Copenhagen University Hospital - Rigshospitalet, DK-2600 Glostrup, Denmark; Translational Research Centre, Copenhagen University Hospital – Rigshospitalet, DK-2600 Glostrup, Denmark; RNA Therapeutics, Copenhagen University Hospital – Rigshospitalet, DK-2600 Glostrup, Denmark; Section for Cell and Drug Technologies, Department of Health Technology, Danish Technical University, 2800, Kgs. Lyngby, Denmark; Department of Growth and Reproduction, Copenhagen University Hospital – Rigshospitalet Glostrup, DK-2100 Copenhagen Ø, Denmark; Department of Science and Environment, Roskilde University, DK-4000 Roskilde, Denmark; Department of Anesthesia, Critical Care and Pain Medicine, Beth Israel Deaconess Medical Center, Harvard Medical School, Boston, Massachusetts 02215, USA; Center for Translational Neuromedicine, University of Copenhagen, Copenhagen, Denmark

## Abstract

ATP-sensitive potassium (K_ATP_) channel opener levcromakalim is a potent inducer of vasodilation, headache, and migraine attacks in humans and tactile hypersensitivity in mice. Other migraine-inducing agents such as nitric oxide (NO) donors, CGRP, and PACAP are thought to activate second messengers leading to K_ATP_ opening. Yet, how K_ATP_ channel opening leads to migraine remains unclear. Here, we investigated the contribution of nitric oxide synthase (NOS) isoforms and downstream signaling cascades in a mouse model of migraine-relevant tactile hypersensitivity induced by repeated administration of levcromakalim. The non-selective NOS inhibitor N^G^-nitro-L-arginine methyl ester (L-NAME) effectively prevented levcromakalim-induced hypersensitivity. Gene expression analysis in the dura mater suggested contributions from endothelial NOS (eNOS) and inducible NOS (iNOS). Semi-selective nNOS inhibition with S-methyl-L-Thiocitrulline (SMTC) or genetic deletion of neuronal NOS (nNOS) had minimal effect on hypersensitivity and no effect on vasodilation. In contrast, *eNOS^-/-^* mice were partially protected from levcromakalim-induced hypersensitivity and exhibited impaired vascular response, highlighting eNOS as a key mediator. Inhibition of iNOS with *S-*methylisothiourea (SMT) revealed a possible contribution from iNOS as well. Surprisingly, inhibition of soluble guanylate cyclase (sGC) had no effect, while the peroxynitrite decomposition catalyst FeTPPS partially attenuated hypersensitivity, implicating nitrosative stress—rather than classical NO-sGC-cGMP signaling—as the critical downstream pathway. We propose that levcromakalim induces both coupled and uncoupled eNOS activity, enhanced NO production and generation of reactive nitrogen species, including peroxynitrite. Our findings reveal a pivotal role for eNOS and peroxynitrite in K_ATP_ channel-induced migraine-relevant hypersensitivity and support targeting nitrosative stress as a potential therapeutic strategy.

## Introduction

Migraine is a major public health concern with 1 billion affected people world-wide [61]. It is a primary headache disorder with considerable societal and private losses due to its high disability burden, especially in women under 50 years of age [61,62,69]. Significant therapeutic advances have been made in recent years with the development and marketing of monoclonal antibodies and small molecule receptor antagonists targeting calcitonin gene-related peptide (CGRP) or its receptor [26]. Unfortunately, far from all patients benefit from these drugs, and there is a need to identify alternative or supplemental treatment targets [24].

Experimentally, headache, migraine attacks, and cluster headache can be induced by a series of vasodilating substances including the ATP-sensitive potassium (K_ATP_) channel opener levcromakalim [5–7]. K_ATP_ channel opening is generally considered a common downstream event of several headache-inducing compounds, such as CGRP, pituitary adenylate cyclase-activating polypeptide (PACAP) and nitric oxide (NO)-donors [4]. However, the molecular mechanisms leading from K_ATP_ channel opening to migraine pain remains unknown. Clues from previous clinical and preclinical studies suggest that the effect is mediated via smooth muscle (vascular) and not neuronal K_ATP_ channels in the dura mater [16,20,43,48].

Major lines of evidence support that NO is clinically relevant in migraine and other headache disorders. First, environmental exposure to NO was linked to headaches in people working with explosives [67]. Second, controlled experimental trials have firmly established an important role of NO in headache as infusion of NO-donor glyceryl trinitrate (GTN) induce migraine, tension type headache, and cluster headache [9,14,39,66]. Third, spontaneous migraine attacks [45] and tension type headaches [10] were aborted by L-N^G^ methylarginine hydrochloride (L-NMMA), a non-selective inhibitor of nitric oxide synthases (NOS). There are three NOS isoforms: neuronal (nNOS), endothelial (eNOS), and inducible (iNOS). While nNOS is mainly expressed in neurons, eNOS is expressed at various sites like platelets, endothelium, and several brain areas. Both nNOS and eNOS are constitutively active while iNOS is induced as part of the immune response system [52]. It is not clear if all or one of the three NOS isoforms are important in migraine, and what the downstream mechanisms of endogenous NO production is in headache pathophysiology.

Here, we investigated NO involvement in response to levcromakalim in mice. *In vivo*, levcromakalim exposure every other day induce hypersensitivity to mechanical and heat stimulation. Also, c-Fos expression in the trigeminal nucleus caudalis (TNC) is amplified in response to lecromakalim exposure [8,21]. Mechanisms were investigated by chemical and genetic approaches *in vivo* and *ex vivo*. The aim of the current study was to investigate the possible involvement of the gaseous transmitter NO in levcromakalim-induced vasodilation and tactile hypersensitivity in mice. We investigate both NO formation via the nitric oxide synthases and subsequent activity of NO.

Our findings suggest a crucial role of eNOS in levcromakalim-mediated arterial dilation and mechanical hypersensitivity leading to subsequent formation of peroxynitrite (PN). These findings are a most relevant step towards improved understanding of the biochemical mechanism of migraine and other headache disorders.

## Materials and methods

### Animals

A total of 473 mice from three different sub strains on C57BL/6 background were used to complete this study: i) wildtype (WT) C57BL/6JBomTac (Taconic, Denmark), N_WT_ =413. ii) B6.129S4-*Nos1^tm1Plh^*/J (*nNOS^-/-^*) mice delivered from Jackson Laboratory (strain #: 002986, JAX, USA), originally developed by Huang et al. (1993) [37], N ^-/-^ =30. Due to time restraints, WT mice from Taconic were used as controls for the *nNOS^-/-^* strain. iii) BKS.129P2(Cg)-*Nos3^tm1Unc^*/J (*eNOS^-/-^*) JAX (strain #: 018295), originally developed by Zhao et al. (2006) [72]. Breeders from JAX were backcrossed to C57BL/6JBomTac in-house to generate matching KO and WT controls, N_eNOS_^-/-^=30. See supplementary materials for genotyping protocol. Female and male adult mice (7-14 weeks old) were used, weighing 18– 35 g. Mice not bred in house were left to acclimatize for at least one week before experiments. Mice were age and sex matched within individual experiments.

Mice were group housed as previously described [28]. The light cycle was 12 h with lights on at 7 am, temperature 21-23°C, relative humidity 55-65%, and free access to food and water. Multiple shelters and types of nesting materials were provided for enrichment.

All experiments and breeding were approved by the Danish Animal Experiments Inspectorate (ethical approval numbers 2019-15-0201-00378, 2022-15-0201-01347, 2017-15-0201-01358) and carried out according to ARRIVE guidelines. All *in vivo* and *ex vivo* experiments are summarized in Table 1.

**Table 1.** – Overview of experiments, test groups, compounds used and number of mice.

|  | Figure | Type of experiment | Treatments (n) | Notes |
| --- | --- | --- | --- | --- |
| Figure 1 | B-D | Von Frey | 1. L-NAME + VEH (LEV) ($n_{WT} = 10$)<br>2. L-NAME + LEV ($n_{WT} = 10$)<br>3. VEH (L-NAME) + LEV ($n_{WT} = 10$) | Tissue taken for qRT-PCR (Fig. 1F-H) |
| | E | Rotarod | 1. MIDA ($n_{WT} = 24$)<br>2. LEV ($n_{WT} = 24$)<br>3. L-NAME ($n_{WT} = 23$) | Mice from another cohort |
| | F-H | qRT-PCR | 1. VEH ($n_{WT} = 12$)<br>2. LEV ($n_{WT} = 12$) | Tissue taken after VF protocol (Fig. 1B-D) |
| Figure 2 | B | Rotarod | 1. VEH (SMTC)+ VEH (LEV) ($n_{WT} = 9$)<br>2. SMTC + VEH (LEV) ($n_{WT} = 9$)<br>3. SMTC + LEV ($n_{WT} = 9$)<br>4. VEH (SMTC) + LEV ($n_{WT} = 9$)<br>1. MIDA ($n_{WT} = 12$) | Mice from VF (Fig. 2C-E) |
| | C-E | Von Frey | 2. VEH (SMTC) + VEH (LEV) ($n_{WT} = 12$)<br>3. SMTC + VEH (LEV) ($n_{WT} = 12$)<br>4. SMTC + LEV ($n_{WT} = 12$)<br>5. VEH (SMTC) + LEV ($n_{WT} = 12$) | Mice also tested on rotarod (Fig. 2B). |
| Figure 3 | C-D, F-G | Von Frey | 1. WT + VEH (LEV) ($n_{WT} = 12$)<br>2. WT + LEV ($n_{WT} = 12$)<br>3. $nNOS^{-/-}$ + VEH (LEV) ($n_{nNOS^{-/-}} = 12$)<br>4. $nNOS^{-/-}$ + LEV ($n_{nNOS^{-/-}} = 12$) | Mice also tested on rotarod (Fig. 3E) |
| | E | Rotarod | 1. WT + VEH (LEV) ($n_{WT} = 12$)<br>2. WT + LEV ($n_{WT} = 12$)<br>3. $nNOS^{-/-}$ + VEH (LEV) ($n_{nNOS^{-/-}} = 12$)<br>4. $nNOS^{-/-}$ + LEV ($n_{nNOS^{-/-}} = 12$) | Mice from VF (Fig. 3C-D + F-G) |
| | H-I | Myograph | 1. WT ($n_{WT} = 6$)<br>2. $nNOS^{-/-}$ ($n_{nNOS^{-/-}} = 6$) | |
| Figure 4 | C-D,<br>F-G | Von Frey | 1. $eNOS^{+/+}$ + VEH (LEV) ($n_{eNOS^{+/+}} = 14$)<br>2. $eNOS^{+/+}$ + LEV ($n_{eNOS^{+/+}} = 16$)<br>3. $eNOS^{-/-}$ + VEH (LEV) ($n_{eNOS^{-/-}} = 16$)<br>4. $eNOS^{-/-}$ + LEV ($n_{eNOS^{-/-}} = 14$) | Tissue taken for myograph (Fig. 4H-J) |
| | E | Rotarod | 1. $eNOS^{+/+}$ + VEH (LEV) ($n_{eNOS^{+/+}} = 5$)<br>2. $eNOS^{+/+}$ + LEV ($n_{eNOS^{+/+}} = 7$)<br>3. $eNOS^{-/-}$ + VEH (LEV) ($n_{eNOS^{-/-}} = 10$)<br>4. $eNOS^{-/-}$ + LEV ($n_{eNOS^{-/-}} = 17$) | Mice from VF (Fig 4C-D + F-G) |
| | H-J | Myograph | 1. $eNOS^{+/+}$ ($n_{eNOS^{+/+}} = 5$)<br>2. $eNOS^{-/-}$ ($n_{eNOS^{-/-}} = 7$) | Tissue taken after VF protocol (Fig 4C-D + F-G) |
| Figure 5 | B-D | Von Frey | 1. SMT + VEH (LEV) ($n_{WT} = 16$)<br>2. SMT + LEV ($n_{WT} = 16$)<br>3. VEH (SMT) + LEV ($n_{WT} = 16$) | Mice also tested on rotarod (Fig. 5E) |
| | E | Rotarod | 1. SMT + VEH (LEV) ($n_{WT} = 16$)<br>2. SMT + LEV ($n_{WT} = 16$)<br>3. VEH (SMT) + LEV ($n_{WT} = 16$) | Mice from VF (Fig. 5B-D) |
| | F-H | Myograph | 1. SMT ($n_{WT} = 6$)<br>2. VEH ($n_{WT} = 6$) | Tissue taken after VF protocol (Fig 5B-D) |
| Figure 6 | B-D | Von Frey | 1. VEH (ODQ) + VEH (LEV) ($n_{WT} = 12$)<br>2. ODQ + VEH (LEV) ($n_{WT} = 12$)<br>3. ODQ + LEV ($n_{WT} = 12$)<br>4. VEH (ODQ) + LEV ($n_{WT} = 12$) | - |
| | E-G | Von Frey | 1. ODQ + VEH (LEV) ($n_{WT} = 12$)<br>2. ODQ + LEV ($n_{WT} = 12$)<br>3. VEH (ODQ) + LEV ($n_{WT} = 12$) | - |
| Figure 7 | B-D | Von Frey | 1. FeTPPS + VEH (LEV) ($n_{WT} = 12$)<br>2. FeTPPS + LEV ($n_{WT} = 12$)<br>3. VEH (FeTPPS) + LEV ($n_{WT} = 12$) | Mice also tested on rotarod (Fig. 7E). |
| | E | Rotarod | 1. FeTPPS + VEH (LEV) ($n_{WT} = 9$)<br>2. FeTPPS + LEV ($n_{WT} = 9$) | Mice from VF (Fig. 7B-D) |
| | | | 3. VEH (FeTPPS) + LEV ($n_{WT} = 9$)<br>4. MIDA ($n_{WT} = 9$) | |
| | F-I | Western blot | 1. VEH ($n_{WT} = 6$)<br>2. LEV ($n_{WT} = 6$) | Tissue also used<br>for qRT-PCR<br>(Fig. 7J-K) |
| | J-K | qRT-PCR | 1. VEH ($n_{WT} = 12$)<br>2. LEV ($n_{WT} = 12$) | Tissue also used<br>for WB (Fig. 7F-I) |

L-NAME = N<sup>G</sup>-nitro-L-arginine methyl ester, MIDA = midazolam, SMTC = S-methyl-L-Thiocitrulline, SMT = S-methylisothiurea, ODQ = 1H-[1,2,4]oxadiazolo[4,3-a]quinoxalin-1-one, FeTPPS = Fe(III)5,10,15,20-tetrakis(4-sulfonatophenyl)porphyrinato chloride, VF = von Frey

### The Levcromakalim Mouse Model of Provoked Migraine

We applied the mouse model of levcromakalim-induced migraine with hind paw cutaneous tactile sensitivity as measuring endpoint [17,20,27,28,34,53]. Levcromakalim induce a global hypersensitivity that can be detected both in cephalic and hind paw dermatomes [20].

Mice were sensitized by repeated injections of levcromakalim 1 mg/kg i.p. [19–21]. Chemical inhibitors were injected 15-20 min prior to levcromakalim. Each test paradigm lasted nine days with injections and von Frey sensitivity testing every other day (days 1, 3, 5, 7, and 9)). Mice were habituated to test chambers 45 min prior to the first test day and every test day 30-45 min before each measurement. On test days, two von Frey measurements were made: a) basal values prior to injections and b) 2 h post injection of levcromakalim. Animals were allocated to test groups after the first baseline measurement on day 1. Tactile sensitivity was measured on the left hind paw with nylon von Frey (VF) filaments (0.008 – 2.0 g, excluding 1.4 g, Ugo Basile, Italy) using the up-down method [15]. Subsequently, 50% withdrawal thresholds were calculated using the open access online calculator optimized and described by Christensen et al. 2020 [18]. Exact delta values and filament force were applied as settings. Experimenters were blinded to treatment groups.

### Motor function (Rotarod)

General motor function was examined to ensure that potential side effects of the study drugs, such as impaired motor function or sedation, did not bias the cutaneous sensitivity test with VF filaments. Two different rotarods and protocols were applied due to unavoidable change of equipment. The first experiment (L-NAME) was performed on a rotarod (LE820 Panlab, Harvard apparatus) and the other experiments were performed on Rotarods Advanced; (IITC Life Science). Motor function was assessed immediately following the VF tests on test day 7 or 9 corresponding to 2.5-3 h after levcromakalim administration. For assessment of L-NAME (Fig 1E) only a single dose was given in a group of mice not included in VF experiments. No observations or later tests indicated that results would have been different if the mice had been subjected to a full VF protocol. In the LE820 Panlab protocol (Fig 1E), mice were trained to stay on the rotarod for two min at 15 rpm prior to drug challenge. Following drug administration, each mouse was given three attempts on the rotarod and the longest duration on the rotarod was recorded. For the Rotarods Advanced protocol (Fig 2B, 3E, 4E, 5E, 7E) each mouse was given a single attempt on the rotarod, starting from 0 rpm and gradually accelerating to 30 rpm within 45 seconds, with a maximum duration of 150 seconds. The duration spent on the rotarod was recorded. Midazolam (2 mg/kg, intraperitoneally (i.p.)) was administered as positive control, 10-25 min prior to testing. Mice were randomly allocated into treatment groups when receiving midazolam.

**Figure 1:**
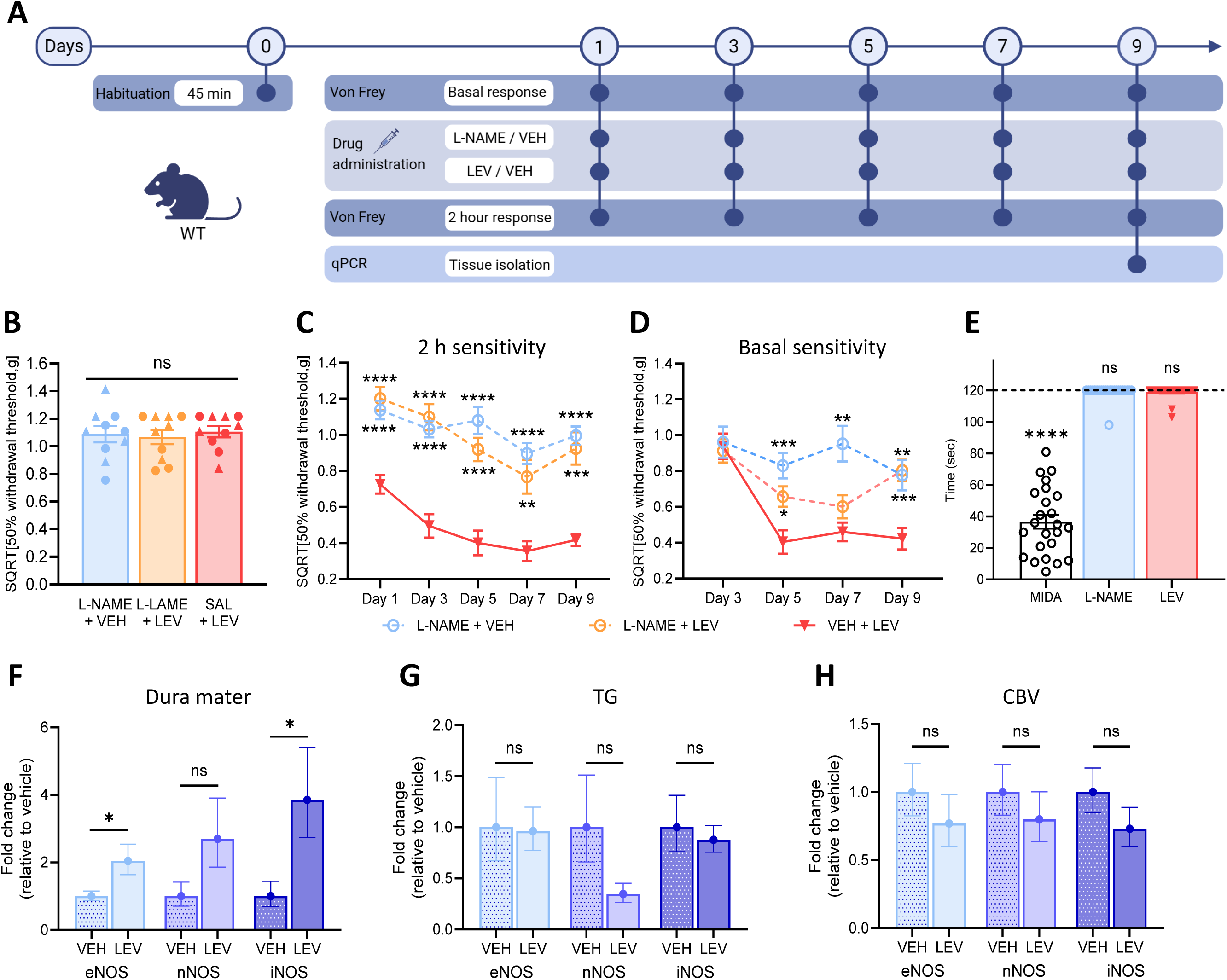
Effect of NOS inhibition by L-NAME on levcromakalim-induced hypersensitivity. A) Design and timeline of the test paradigm for levcromakalim (LEV)-induced hypersensitivity and NG-nitro-l-arginine methyl ester hydrochloride (L-NAME) administration. One day before drug administration, WT mice were placed in the test chambers for 45 min (Day 0 – Habituation). Cutaneous tactile sensitivity measurements on the plantar surface of the left hind paw were performed using von Frey (VF) filaments on every other day before (Basal response) and 2 hours (2 h response) after drug administration for a total of 9 days. Before each VF test, mice were placed in test chambers 30-45 min prior to testing. On every test day, animals received L-NAME (100 mg/kg i.p.) 15-20 minutes prior to LEV (1 mg/kg i.p.) or vehicle (2% DMSO in saline) administration. Following the last VF test on day 9, the dura mater, trigeminal ganglion (TG) and cerebral blood vessels (CBV) were isolated for quantitative real-time PCR measurements. The illustration is created with Biorender.com. B) Baseline tactile sensitivity values on day 1. Data are presented as individual data points with triangles representing males (▲) and circles representing females (●). One-way ANOVA with *post hoc* Tukey multiple comparison test. C-D) Tactile sensitivity measurements day 1-9 (basal response day 1 is shown in B). Sensitivity was measured C) acutely 2 hrs. post LEV or VEH administration and D) prior to drug injections. Data are presented means ± SEMs and calculated as 50% withdrawal threshold (g) after square root transformation (SQRT). Mixed-effects analysis with Dunnett’s correction for multiple comparison (* = compared to positive control (VEH+LEV)). E) Motor function was examined after single injection of LEV, L-NAME, and midazolam administration (MIDA, 2 mg/kg). Data are shown as individual data points with mean ± SEM. Wilcoxon signed-rank test comparing to the hypothetical value of 120. F-H) mRNA expression of *Nos3*, *Nos1* and *Nos2* (encoding eNOS, nNOS and iNOS, respectively) in F) dura mater, G) TG and H) CBV from mice treated with LEV or vehicle. Gene expression was normalized to *Hrpt*. Data are presented as mean ± SEM. Multiple unpaired t-tests with Holm-Šídák correction for multiple comparison. Dura mater: eNOS *P* = 0.0469, nNOS *P* = 0.0794, iNOS *P* = 0.0469. TG: eNOS *P* = 0.939, nNOS *P* = 0.152, iNOS *P* = 0.903. CBV: eNOS *P* = 0.696, nNOS *P* = 0. 696, iNOS *P* = 0.567, * *= P* < 0.05, ** *= P* < 0.01, *** *= P* < 0.001, **** *= P* < 0.0001.

**Figure 2:**
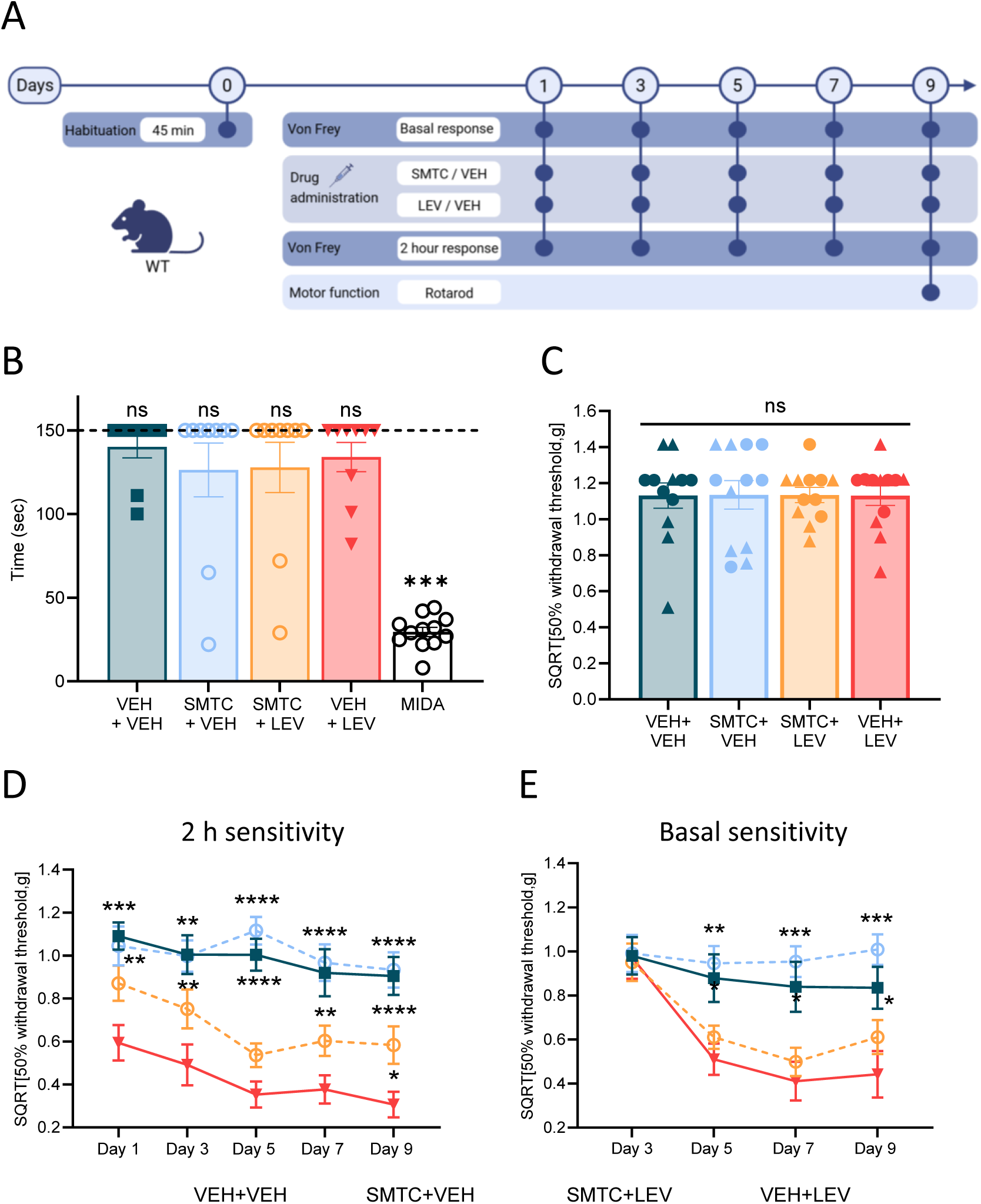
Effect of chemical inhibition of nNOS by SMTC on levcromakalim-induced hypersensitivity. A) Design and timeline of the test paradigm for levcromakalim (LEV)-induced hypersensitivity and S-methyl-L-thiocitrulline (SMTC) administration. One day before drug administration, WT mice were placed in the test chambers for 45 min (Day 0 – Habituation). Cutaneous tactile sensitivity measurements on the plantar surface of the left hind paw were performed using von Frey (VF) filaments on every other day before (Basal response) and 2 hours (2 h response) after drug administration for a total of 9 days. Before each VF test, mice were placed in test chambers 30-45 min prior to testing. On every test day, animals received SMTC (60 mg/kg i.p.) 15-20 minutes prior to LEV (1 mg/kg i.p.) administration. Following the last VF test on day 9, motor function after drug administration was testing using a rotarod with midazolam (MIDA, 2 mg/kg i.p.) as a positive control. The illustration is created with Biorender.com. B) Motor function measured by rotarod performance on day 9 and after midazolam administration. Data are shown as individual data points with mean ± SEM. Wilcoxon signed-rank test comparing to the hypothetical value of 150 (max duration of the test). C) Baseline tactile sensitivity values on day 1. Data are presented as individual data points with triangles representing males (▲) and circles representing females (●). One-way ANOVA with *post hoc* Tukey multiple comparison test. D-E) Tactile sensitivity measurements day 1-9 (basal response day 1 is shown in C). Sensitivity was measured D) acutely 2 hrs. post LEV or VEH administration and E) prior to drug injections. Data are presented means ± SEMs and calculated as 50% withdrawal threshold (g) after square root transformation (SQRT). Mixed-effects analysis with Dunnett’s correction for multiple comparison (* = compared to positive control (VEH+LEV)). * *= P* < 0.05, ** *= P* < 0.01, *** *= P* < 0.001, **** *= P* < 0.0001.

### Wire myography

For tissue extraction and wire myography, the same protocol was used as previously described in detail by Ernstsen et al. (2022) [28]. All three strains of mice (WT, *nNOS^-/-^*, and *eNOS^-/-^*) were euthanized without damaging the carotid artery (100 µL pentobarbital i.p. (200 mg/ml pentobarbital + 20 mg/ml lidocaine, Hospital Pharmacy, Copenhagen, Denmark)). Briefly, the right common carotid artery was dissected and cut into segments of approximately 1 mm, immersed in freshly made, oxygenated Na^+^ Krebs buffer and mounted on wires (40 µm) in a Mulvany-Halpern wire myograph (Danish Myo Technology, Denmark) held at 37°C. After 15 min equilibration, contractility of tissue was examined by K^+^ to validate functionality of arteries. Precontraction was induced by U46619 (0.1 µM) before the effects of levcromakalim (0.1 µM, 1 µM, 10 µM and 30 µM) or vehicle (0.001 – 0.3% DMSO) on the vessels were examined. For *eNOS^-/-^* mice, the segments were precontracted again to functionally test eNOS deletion with carbachol (0.01 µM, 0.1 µM, 1 µM and 10 µM). Buffer was continuously changed during experiments and relaxation was calculated as percentage relative change in vessel tension within individual samples using the formula below (1) [28].

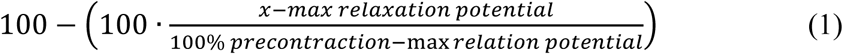

where x is the tension of the blood vessel after stimulation. 100% precontraction and max relaxation potential are constants for individual blood vessels. Each data point represents relaxation of either one segment or mean of two segments.

### Quantitative real-time mRNA analysis using polymerase chain reaction (qRT-PCR)

Quantitative real-time PCR analysis of target mRNA levels mere made on trigeminal ganglion (TG), cerebral blood vessels (CBV), and dura mater. Tissue was isolated immediately after anesthesia with gas (70% CO_2_ and 30% O_2_) and euthanasia by decapitation and kept on dry ice until stored at −80°C. RNA was extracted using a combined homogenization and trizol-based approach (See supplementary material). 200 ng total RNA was reversed transcribed using the iScript cDNA Synthesis Kit (Bio-Rad, USA) according to the manufacturer’s protocol. qRT-PCR was performed either with SYBR green (Fig 1) or TaqMan-based (Fig 7) detection methods. SYBR green-based qRT-PCR reactions were performed using RNase free water, 2x pre-designed SensiFast mix (Meridan Bioscience, USA), forward and reverse primers (table 2) and cDNA. The thermal cycling condition included an initial step at 95 °C for 2 min followed by 40 PCR cycles at 95 °C for 5 s and 60 °C for 30 s. TaqMan-based qRT-PCR reactions were performed in a 10μl reaction volume containing RNase free water, 20x pre-designed TaqMan mouse specific gene expression assay (IDT, USA), PrimeTime Gene expression Master Mix (IDT, USA), and 2μl cDNA. The thermal cycling condition included an initial denaturation step at 95 °C for 2 min followed by 40 PCR cycles at 95 °C for 5 s and 60 °C for 30s, and lastly a melt curve program of 95C for 5s, 60C for 30 s., and 95C for 5 s. All qRT-PCR reactions were performed using the Quant-Studio 6 Pro Real-Time PCR system (Applied Biosystems, USA). See pre-designed TaqMan gene expression assay used in Table 2.

**Table 2.**
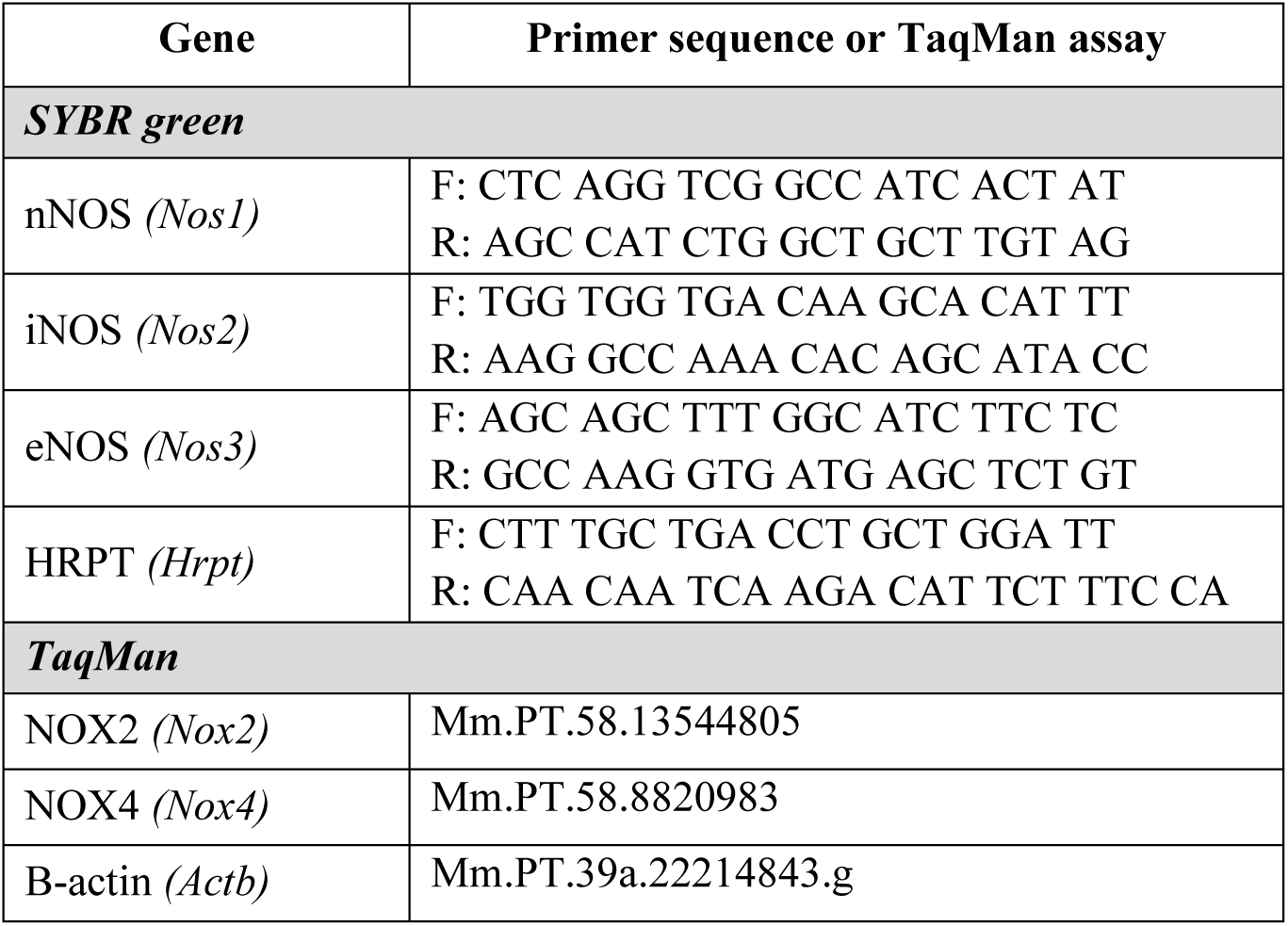
– List of primer sequence used and TaqMan assay. Primers were ordered from TAG Copenhagen.

### Western blot

Following RNA isolation, the trizol supernatant were kept for protein isolation (see supplementary material). Protein concentration was determined using a Bio-Rad DC protein assay (Bio-Rad, USA) according to the manufacturer’s manual. 15 mg of TG protein and 13 mg of dura mater protein was loaded to a 4-12% Bis-Tris gel and were transferred to a PVDF membrane using iBlot 2 (Life Technologies, USA). Membranes were blocked with 5% nonfat dry milk in 1× TBST at room temperature for 1 h. The membrane was cut to at the 80 kDa mark to enable incubation with anti-3-nitrotyrosine antibody at 1:3,000 (#ab61392, Abcam) and anti-vinculin (#9264, Merck) at 1:10,000 overnight at 4°C. The following day, membranes were washed 4 times in 1× TBST for 5 min and then incubated with 1:3,000 anti-mouse secondary antibody at room temperature for 1 h. Subsequently, membranes were washed in 1× TTBS 4 times for 5 min, and signals were detected using ECL Prime Western Blotting Detection Reagent (Cytiva, USA). Protein bands were visualized using Las-4000 (FujiFilm, Japan). The quantification analysis was performed using ImageJ and normalized to vinculin expression.

### Study compounds

Compounds used *in vivo* and *ex vivo* are listed in Table 3 with all relevant information regarding dose, solvent, and vehicle. All compounds were injected i.p. in the lower right side of the mouse in a volume of 10 ml/kg.

**Table 3.** – List of compounds used in *in vivo* and *ex vivo* studies.

| Compound | Mechanism of Action | Solvent | Dose | Vehicle |
| --- | --- | --- | --- | --- |
| ***In vivo*** |  |  |  |  |
| Levcromakalim | K<sub>ATP</sub> channel opener | 2% DMSO in Saline | 1 mg/kg, i.p. | 2% DMSO in Saline |
| L-NAME | Non-selective NOS inhibitor | Saline | 100 mg/kg i.p. | Saline |
| SMTC | Semi-selective nNOS inhibitor | DPBS | 60 mg/kg i.p. | DPBS |
| SMT | Selective iNOS inhibitor | Saline | 10 mg/kg i.p. | Saline |
| ODQ | sGC inhibitor | 2% DMSO in Saline | 1 mg/kg i.p.<br>10 mg/kg i.p. | 2% DMSO in Saline |
| FeTPPS | Peroxynitrite decomposition catalyst | Saline | 30 mg/kg i.p. | Saline |
| Midazolam | Benzodiazepine | Saline | 2mg/kg i.p. | - |
| ***Ex vivo*** |  |  |  |  |
| Levcromakalim | K<sub>ATP</sub> channel opener | DMSO in Na<sup>+</sup> Krebs | 0.1 µM, 1 µM, 10 µM, 30 µM |  |
| Carbachol | Acetylcholine receptor agonist | Na<sup>+</sup> Krebs | 0.01 µM, 0.1 µM, 1 µM, 10 µM, |  |

DMSO = dimethyl sulfoxide. L-NAME = N<sup>G</sup>-nitro-L-arginine methyl ester. SMTC = S-methyl-L-Thiocitrulline. SMT = S-methylisothiurea. sGC = soluble guanylyl cyclase. FeTPPS = 5,10,15,2 – Tetrakis(4-sulfonatophenyl)porphyrinato-ferro (III) chloride. DPBS = Dulbecco's phosphate-buffered saline.

### Statistical Analyses

All data analyses and visualization were made in GraphPad Prism 10.4.1. Statistical summary data for all data are presented in Table 4.

**Table 4.**
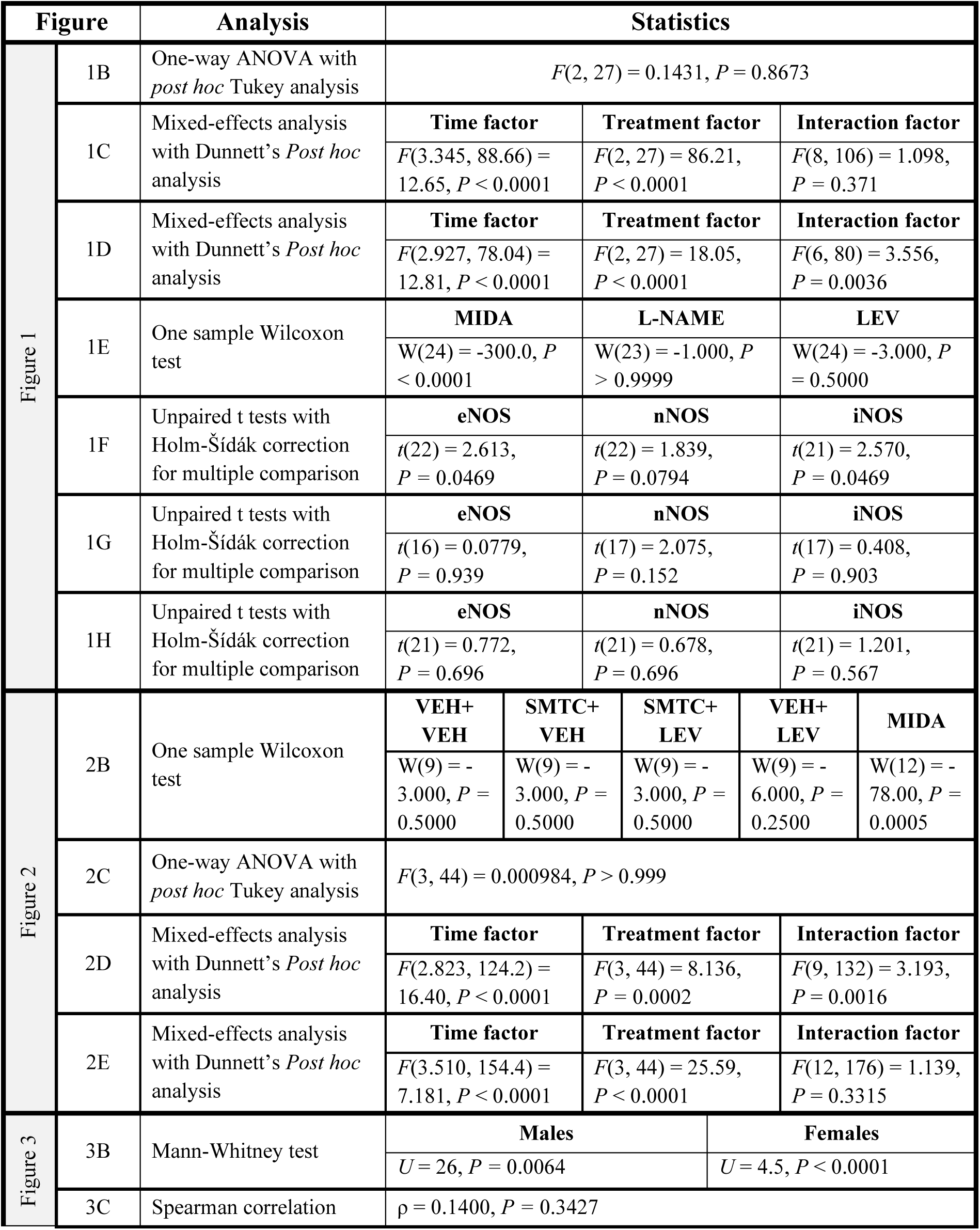
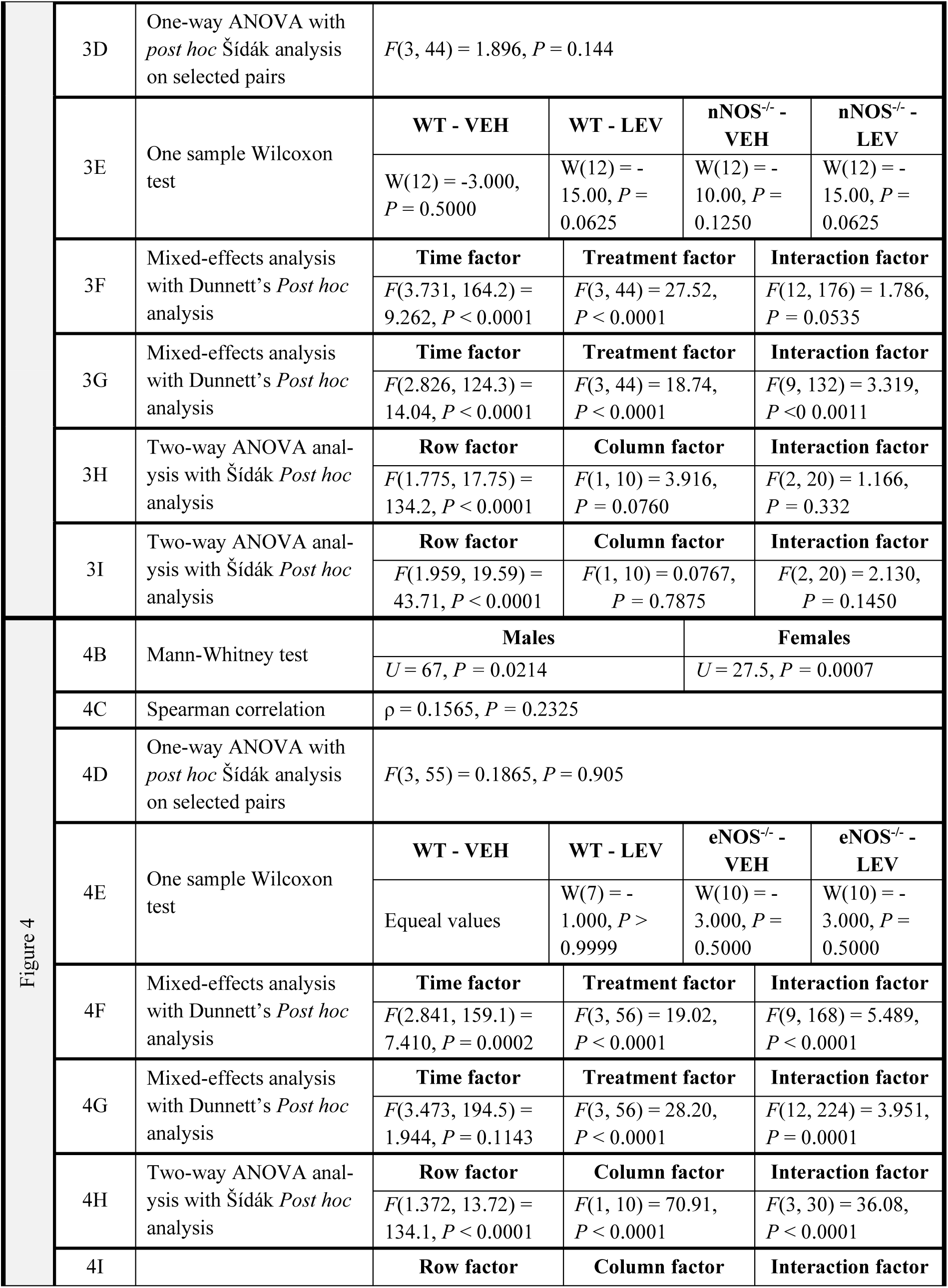
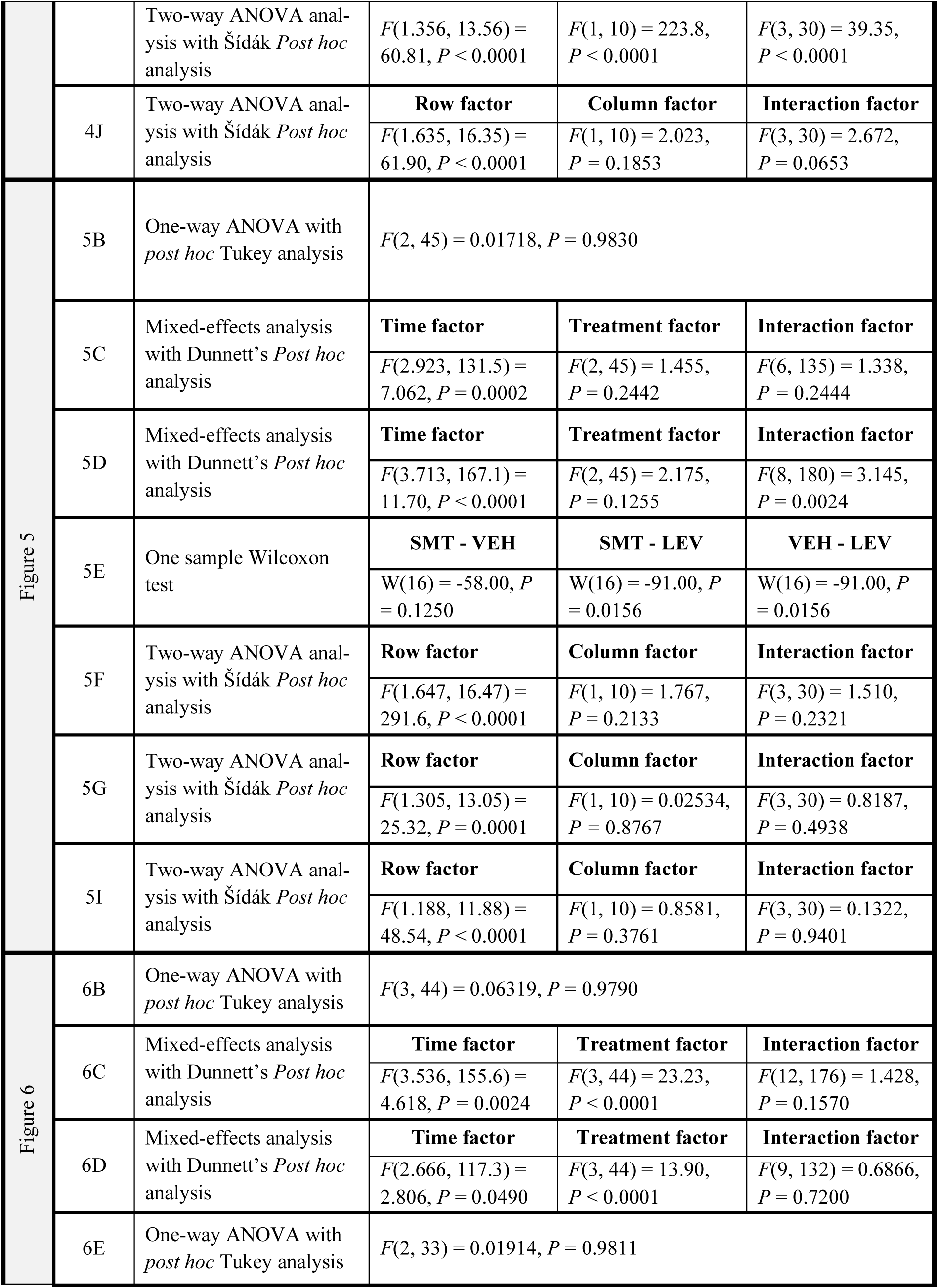
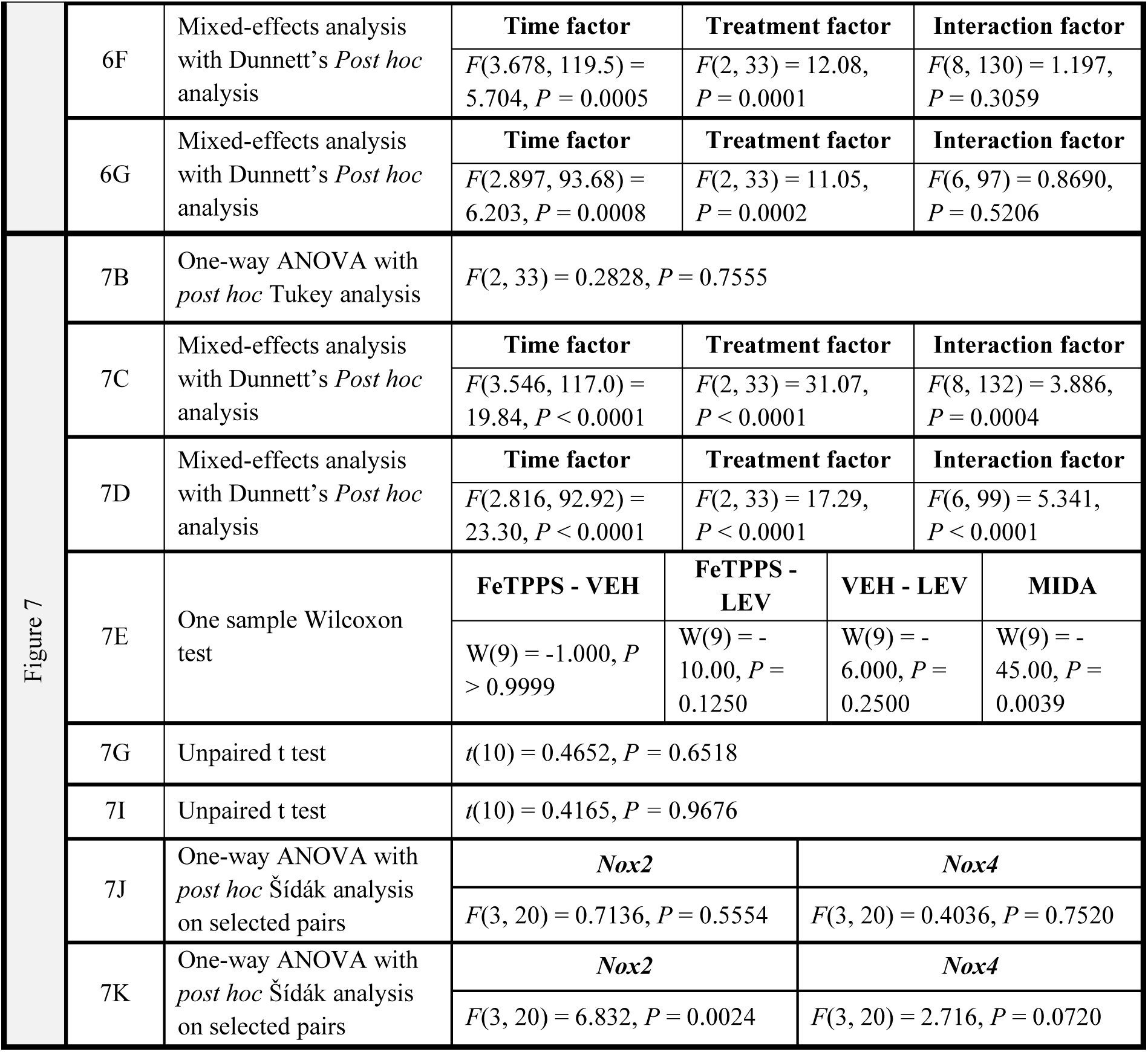
- Statistical summary data for all data.

#### Von Frey data

Mice were allocated to treatment groups with stratification according to baseline 50 % withdrawal thresholds, sex, and home cage. 50% withdrawal thresholds were square root transformed for improved gaussian distribution. Data is presented as mean ± SEM in text and graphs. Baseline values were compared by one-way ANOVA with Tukey’s post-hoc test assuming equal variances. Repeated test paradigms were analyzed by mixed-effect analysis with Dunnet’s post hoc comparison to the positive control group (VEH-LEV). Sphericity was not assumed, so Geisser-Greenhouse correction was applied.

##### Body weights

Some groups did not follow a gaussian distribution, and therefore weights were compared using the Mann-Whitney test. Spearman correlations were calculated to assess if differences in body weight between *eNOS^-/-^* and *nNOS^-/-^* mice compared to WT affected the 50% withdrawal thresholds.

#### Rotarod

Wilcoxon signed Rank Test with the hypothetical value of 150 as this is the maximal duration mice were allowed on the rotarod (In Fig 1E, the hypothetical value was 120 due to alternative test protocol). Data are presented as mean ± SEM in text and graph as well as individual points.

#### Wire myography

Two-way ANOVA with Šídák *Post hoc* analysis was used to compare *nNOS^-/-^* and *eNOS^-/-^* with WT mice within each drug concentration. Data are presented as mean ± SEM in text and graph as well as individual points.

#### qRT-PCR

In Fig 1, multiple unpaired t-tests with Holm-Šídák correction for multiple comparison was used to compare vehicle to levcromakalim exposure. In Fig 7, one-way ANOVA with Šídák *Post hoc* analysis was used to compare vehicle and levcromakalim exposure within sexes and vehicle exposure between sexes. Data are presented as mean ± SEM.

#### Western blot

Unpaired t tests were used to compare vehicle and levcromakalim treatment within each sex. Data are presented as mean ± SEM with individual points.

#### Sample size determination

No a priori sample size calculations were performed for any experiments as effects sizes were not known. Sample sizes were based on our in-house extensive experience with both the *in vivo* model and wire myography technique. For VF experiments initial experiments were performed with n=12. In experiments where animals were lost due to for instance cages with male aggression or if effect sizes were small, a replication experiment was performed and data pooled.

## Results

### NOS inhibition prevented development of levcromakalim-induced hypersensitivity

To investigate if NOS activity is involved in hypersensitivity developed following repeated levcromakalim injections, the non-selective NOS inhibitor N^G^-nitro-L-arginine methyl ester hydrochloride (L-NAME) was injected prior to levcromakalim on every test day. Withdrawal thresholds were measured before and 2 h after levcromakalim administration (Fig 1A). L-NAME effectively blocked hypersensitivity both at the basal and 2 h measurements (Fig 1B-D). The latter was blocked completely, whereas the former was partially inhibited likely reflecting decreasing plasma concentrations of L-NAME at this time point. At baseline day 1, all treatment groups had equal 50% withdrawal thresholds with means ± SEMs as follows: L-NAME+VEH (1.09 ± 0.056), L-NAME+LEV (1.1 ± 0.052) and SAL+LEV (1.11 ± 0.040) (P = 0.87) (Fig 1B). After 2 h, there was an overall significant difference between groups (P < 0.0001) with *post hoc* analysis showing that the mice had acutely developed pronounced hypersensitivity to VF stimulation after the first levcromakalim injection on day 1. Here, mean 50 % withdrawal threshold was 1.14 ± 0.052 in the vehicle (VEH+L-NAME, negative control) group vs 0.73 ± 0.052 in the levcromakalim (VEH+LEV, positive control) group, P < 0.0001. By the 5^th^ injection (day 9), 50% thresholds had dropped to 1.00 ± 0.051 and 0.42 ± 0.034, respectively (P < 0.0001). The L-NAME+LEV group had corresponding values of 1.20 ± 0.065 (P < 0.0001) and 0.92 ± 0.089 (P = 0.0004) on day 1 and 9, respectively (Fig 1C). Basal hypersensitivity was present from day 5 (after two LEV injections) and onwards with an overall significant difference between groups (P < 0.0001). On day 5, the levcromakalim (VEH+LEV) had reduced 50 % withdrawal threshold to 0.40 ± 0.066 compared to 0.83 ± 0.071 in the negative control group (L-NAME+VEH) (P = 0.0007) (Fig 1D). On day 5, L-NAME had a slight impact on hypersensitivity 48 hours after its last injection, overall (0.66 ± 0.057, P = 0.02) (Fig 1D). A VEH+VEH group was not included as we have previously published, that L-NAME alone had no effect on VF thresholds [29]. Importantly, neither levcromakalim (*P* = 0.5000) nor L-NAME (*P >* 0.9999) impaired motor function of the mice (Fig 1E).

Having observed an effect from non-specific NOS inhibition, we tested if expression patterns of NOS enzymes changed after levcromakalim administration (Fig 1F-H). TG, dura mater, and CBV were isolated from mice after the VF protocol day 9. In dura mater, gene expression of *Nos3* (encoding eNOS) (*P =* 0.047) and *Nos2* (encoding iNOS) (*P =* 0.047) was significantly changed, while *Nos1* (encoding nNOS) (*P* = 0.079) showed a non-significant trend towards upregulation after repeated levcromakalim administration (Fig 1F). In TG and CBV, levcromakalim did not change the expression of the NOS enzymes (Fig 1G-H).

### A semi-selective nNOS > eNOS inhibitor minimally attenuated levcromakalim-induced hypersensitivity

Although nNOS expression in dura mater was not found to be changed according to statistical testing, we wanted to investigate if nNOS was involved in the hypersensitivity response caused by levcromkalim. Therefore, we tested the (semi-)selective nNOS inhibitor S-methyl-L-Thiocitrulline (SMTC, 60 mg/kg) (Fig 2A). SMTC is mostly selective for nNOS (Ki_human_ values of 1.2, 11, and 40 nM for nNOS, eNOS, and iNOS, respectively) [30], but how this translates to *in vivo* exposure in mice is uncertain. The dose was selected based on efficacy in other studies [29,41,42]. SMTC had no effect on locomotion, as all test groups had normal locomotor activity examined by rotarod performance (*P* values > 0.2500, MIDA; *P* = 0.0005) (Fig 2B). At baseline, all treatment groups had equal 50% withdrawal thresholds: VEH+VEH (1.13 ± 0.071), SMTC+VEH (1.14 ± 0.079), SMTC+LEV (1.13 ± 0.042) and VEH+LEV (1.13 ± 0.055) (P > 0.9999) (Fig 2C). Levcromakalim significantly induced hypersensitivity at 2 h on all test days compared to the negative control (VEH+VEH) (P ranging from <0.0001 to 0.0021) (Fig 2D). SMTC partially inhibited the effect of levcromakalim but a significant difference between the treatment group (SMTC+LEV) and the positive control (VEH+LEV) was only observed on day 9 with 50% withdrawal thresholds of 0.58 ± 0.087 and 0.31 ± 0.059, respectively (P = 0.04) (Fig 2D). At basal measurements, levcromakalim induced a significant hypersensitivity from day 5 and onwards compared to both negative controls (P value ranging from 0.0002 to 0.03) (Fig 2D). This effect was not prevented by SMTC (Fig 2E).

### Global knockout of nNOS activity did not influence tactile hypersensitivity or vasodilatory responses to levcromakalim

Data from the SMTC study did not demonstrate a robust inhibitory effect. The modest inhibitory effect observed may be attributed to the fact that SMTC is only semi-selective, potentially affecting eNOS in addition to nNOS [30]. Consequently, the observed effect might not reflect true nNOS involvement, but rather partial eNOS inhibition. To clarify the specific role of nNOS, we tested levcromakalim in genetically modified mice not expressing nNOS (Fig 3A). The *nNOS^-/-^* mice had a slightly lower body weight than the WT controls (Fig 3B). Mann-Whitney tests showed a significant difference in body weights between male *nNOS^-/-^* and male WT (P = 0.006) and female *nNOS^-/-^*and female WT (P < 0.0001). *nNOS^-/-^* mice were either 77 or 84 days old at day 1, while WT controls were 69-75 days old. Lower body weights were therefore not due to age but rather a developmental difference caused to difference in genetics. Notably, there was no correlation found between mouse body weight basal 50% withdrawal threshold (Fig 3C), emphasizing that lower body weights had no effect on test results (Spearman’s ρ = 0.1400, *P =* 0.3427). Correspondingly, there was also no difference in basal response to tactile stimulation between *nNOS^-/-^* mice and the WT mice (Fig 3D). The *nNOS^-/-^* mice appeared healthy in terms of general wellbeing and motor function measured on day 7 of VF paradigm (All *P* values >0.0625) (Fig 3E). The difference in weight was not sufficient to impair blinding of the experimenter. Genetic deletion of nNOS did not protect mice from developing hypersensitivity in response to levcromakalim treatment, neither at acute nor basal measuring points (Fig 3F and G). Acutely, there was an overall difference between groups (P < 0.0001) with 2h hypersensitivity present from day 5 (P from < 0.0001 to 0.002). On day 9 there was a small difference between the *nNOS^-/-^* and WT responses to levcromakalim (P = 0.02) (Fig 3F). At basal levels, an overall significant difference between groups was present too (P < 0.0001) with basal hypersensitivity established in WT from day 5 P varying from <0.0001 to 0.02) (Fig 3G). A small difference between genotypes was seen on day 7, where levcromakalim-treated *nNOS^-/-^* mice had higher thresholds than levcromakalim-treated WT mice (P = 0.046).

**Figure 3:**
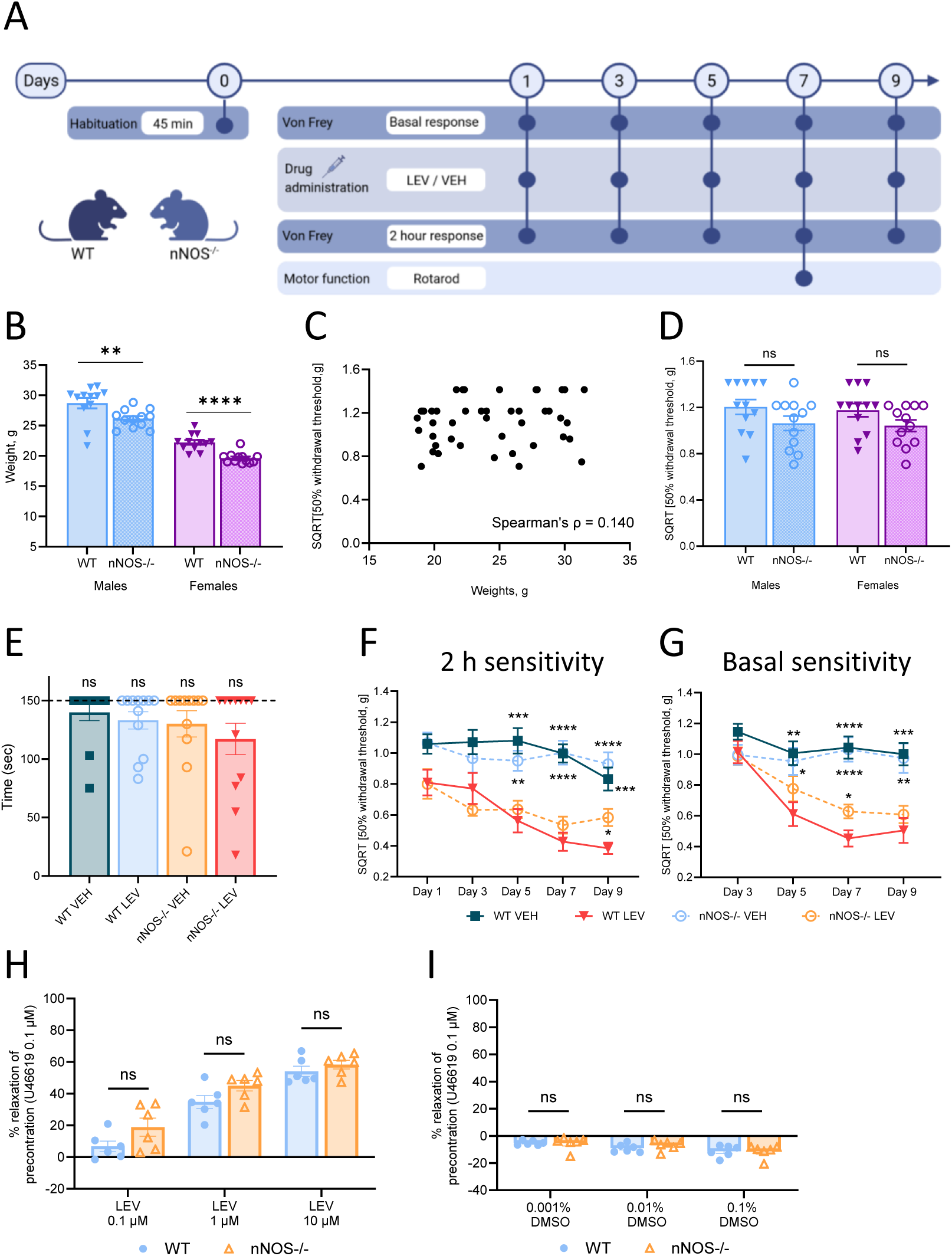
Effect of genetic deletion of nNOS on levcromakalim-induced hypersensitivity using *nNOS^-/-^* mice. A) Design and timeline of the test paradigm for levcromakalim (LEV)-induced hypersensitivity in WT and *nNOS^-/-^* mice. One day before drug administration, mice were placed in the test chambers for 45 min (Day 0 – Habituation). Cutaneous tactile sensitivity measurements on the plantar surface of the left hind paw were performed using von Frey (VF) filaments on every other day before (Basal response) and 2 hours (2 h response) after drug administration for a total of 9 days. Before each VF test, mice were placed in test chambers 30-45 min prior to testing. On every test day, animals received LEV (1 mg/kg i.p.) or vehicle (2 % DMSO in saline). Following the VF tests on day 7, motor function was evaluated by rotarod performance. The illustration is created with Biorender.com. B) Weight of WT and *nNOS^-/-^* animals separated by sex. Data are presented as individual data points with mean ± SEM. Mann-Whitney test, comparing WT and KO within each sex. C) Spearman correlation test between weight (g) and baseline 50% withdrawal thresholds (g). D) Baseline tactile sensitivity values day 1 in WT and *nNOS^-/-^* mice. Data are presented as individual 50% withdrawal threshold (g) after square root transformation (SQRT) with mean ± SEM. One-way ANOVA with *post hoc* Šídák analysis for multiple comparison and is comparing WT and *nNOS^-/-^* within each sex. E) Rotarod performance. Data are shown as individual points with means ± SEM. Wilcoxon signed-rank test comparing to the hypothetical value of 150. F-G) Tactile sensitivity measurements day 1-9 (basal response day 1 is shown in D). Sensitivity was measured F) acutely 2 hrs. post LEV or VEH administration and G) prior to drug injections. Data are presented means ± SEMs and calculated as 50% withdrawal threshold (g) after square root transformation (SQRT). Mixed-effects analysis with Dunnett’s correction for multiple comparison (* = compared to positive control (WT LEV)). H-I) Dilation of the common carotid artery shown as percent of precontraction (U46619 0.1 µM) to increasing concentrations of H) Lev (0.1 µM, 1 µM and 10 µM) and I) vehicle (0.001%, 0.01% and 0.1% DMSO) from WT and *nNOS^-/-^* mice. Relaxation is calculated as % of 100 precontraction. Data are presented as individual data points with mean ± SEM. Two-way ANOVA with *post hoc* Šídák analysis for multiple comparison comparing WT and *nNOS^-/-^* within each concentration. * *= P* < 0.05, ** *= P* < 0.01, *** *= P* < 0.001, **** *= P* < 0.0001.

Isolated carotid arteries from *nNOS^⁻/⁻^* mice had the same dilatory responses to increasing concentrations of levcromakalim as arteries from WT animals (P values ranging from 0.22 to 0.70) (Fig 3H). Specifically, at concentrations of 0.1 µM, 1 µM, and 10 µM levcromakalim, carotid artery dilation in *nNOS^⁻/⁻^* mice was 19 ± 5.7%, 45.0 ± 3.3%, and 58.4 ± 2.7%, respectively, compared to 6.8 ± 3.3%, 34.8 ± 4.1%, and 54.1 ± 3.2% in WT animals. Vehicle exposure (0.001%, 0.01% and 0.1% DMSO) caused no dilation of arteries (*P* = 0.7875) (Fig 3I).

### Global knockout of eNOS compromised arterial dilation and mechanical hypersensitivity induced by levcromakalim

Next, the relevance of eNOS activity was investigated in global eNOS knockout and littermate WT mice (Fig 4A). No eNOS specific chemical inhibitor was available. The *eNOS^-/-^* mice have been reported to be smaller than WT littermates and to have slightly elevated blood pressure [60]. In our animal cohort, *eNOS^-/-^* mice of both sexes had also significantly lower body weights than WT (weights from day 1 of the experiment) Mann-Whitney P = 0.02 for males and P = 0.0007 for females (Fig 4B). However, the *eNOS^-/-^* mice were also slightly younger than their WT counterparts which may have contributed to the smaller size. Male *eNOS^-/-^*mice had a median age of 61 days (range: 51–91), compared to 66 days (range: 55–91 days) for their WT littermate controls. Female *eNOS^-/-^* mice had a median age of 55 days (range: 48–74 days), whereas the female WT littermates had a slightly higher median age of 68 days (range: 55–72). Importantly, there was no correlation between body weight and baseline 50 % withdrawal thresholds (Spearman’s ρ = 0.1565, *P =* 0.2325) (Fig 4C). Although slightly smaller, the *eNOS^-/-^* mice appeared healthy, had baseline 50 % withdrawal thresholds equal to WTs (*P* = 0.905) (Fig 4D) and unimpaired motor performance on the rotarod test after the levcromakalim provocation protocol (All *P* values > 0.5000) (Fig 4E). In the behavioral model, eNOS deletion protected mice from levcromakalim-induced hypersensitivity. WT mice developed hypersensitivity 2 h after the second levcromakalim injection (day 3) (Fig 4F). Here, WT mice had 50 % withdrawal thresholds of 0.66 ± 0.075 and progressed to 0.41 ± 0.096 after the fifth injection (day 9). Corresponding vehicle measures (WT – VEH) were 1.0 ± 0.071 on day 3 (P = 0.0024) and 1.1 ± 0.077 on day 9 (P < 0.0001). *eNOS^-/-^* mice did not develop hypersensitivity after levcromakalim injection reaching a minimum withdrawal threshold of 0.82 ± 0.09 on day 9. *eNOS^-/-^* mice were significantly different from the WTs on test days 3, 5, 7, and 9. P = 0.007-0.03. Similarly, WT mice developed basal hypersensitivity (Fig 4G) after two levcromakalim injections (day 5) where 50 % withdrawal thresholds had dropped to 0.72 ± 0.083 and remained at 1.19 ± 0.081 in the vehicle group, P = 0.0009. This progressed to 0.41 ± 0.072 vs 0.95 ± 0.079 the fourth injection (day 9), P < 0.0001. The progressive development of basal hypersensitivity was not observed in the *eNOS^-/-^* group where basal 50 % thresholds remained above 0.9 throughout the protocol, which was significantly different from the WT levcromakalim group on days 7 and 9. P = 0.0008 and 0.0004, respectively. These data suggest that eNOS activity is an important mediator of levcromakalim-induced hypersensitivity.

**Figure 4:**
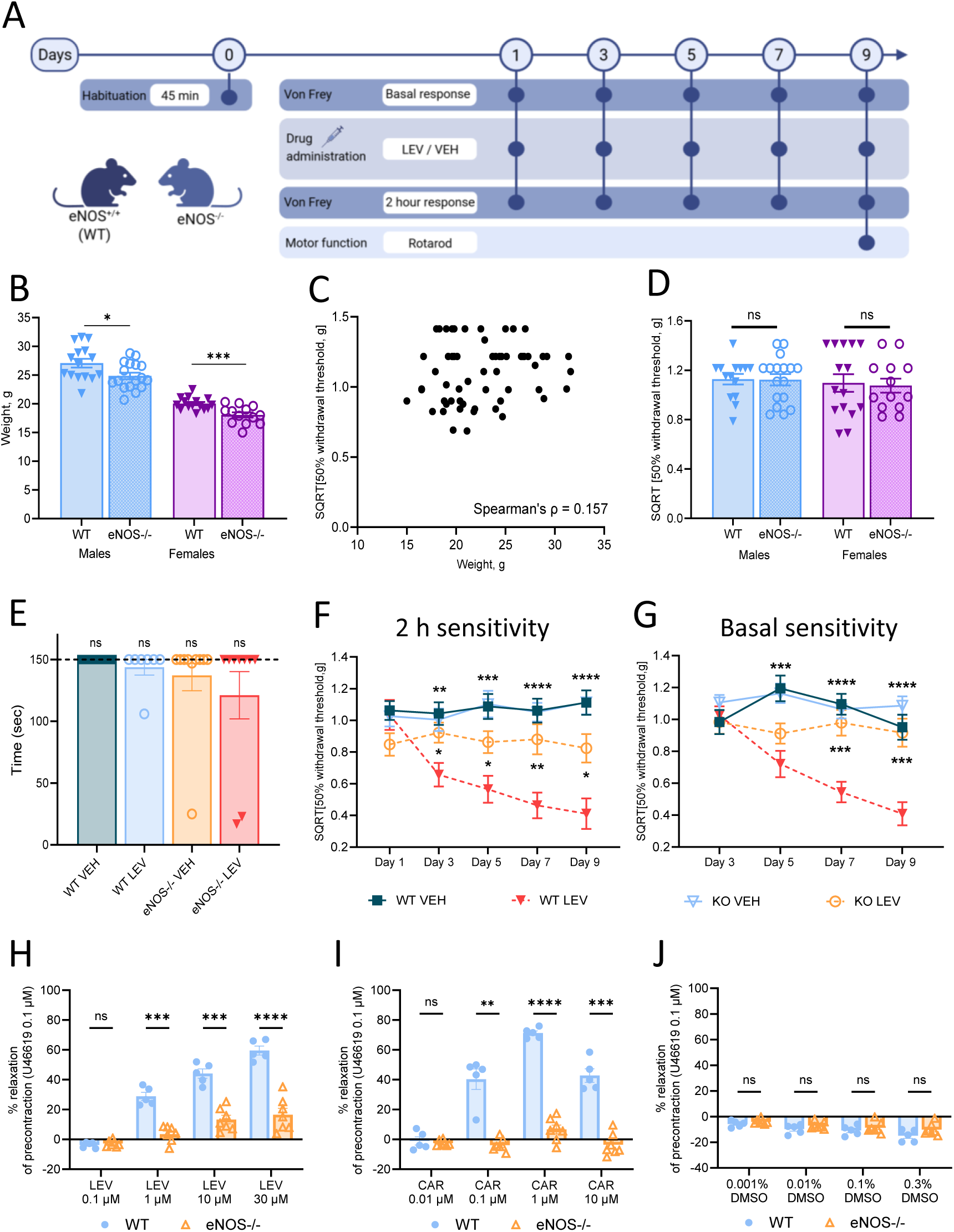
Effect of generic deletion of eNOS on levcromakalim-induced hypersensitivity using e*NOS^-/-^* mice. A) Design and timeline of the test paradigm for levcromakalim (LEV)-induced hypersensitivity in WT and *eNOS^-/-^* mice. One day before drug administration, mice were placed in the test chambers for 45 min (Day 0 – Habituation). Cutaneous tactile sensitivity measurements on the plantar surface of the left hind paw were performed using von Frey (VF) filaments on every other day before (Basal response) and 2 hours (2 h response) after drug administration for a total of 9 days. Before each VF test, mice were placed in test chambers 30-45 min prior to testing. On every test day, animals received LEV (1 mg/kg i.p.) or vehicle (2 % DMSO in saline). Following the final VF test on day 9, motor function was evaluated by rotarod performance. The illustration is created with Biorender.com. B) Weight of WT and *eNOS^-/-^* animals separated by sex. Data are presented as individual data points with mean ± SEM. Mann-Whitney test, comparing WT and KO within each sex. C) Spearman correlation test between weight (g) and basal 50% withdrawal thresholds (g). D) Baseline tactile sensitivity measures on day 1 in WT and *eNOS^-/-^*mice. Data are presented as individual 50% withdrawal threshold (g) after square root transformation (SQRT) with mean ± SEM. One-way ANOVA with *post hoc* Šídák analysis for multiple comparison and is comparing WT and *eNOS^-/-^* within each sex. E) Rotarod performance day 9. Data are shown as individual points with means ± SEM. Wilcoxon signed-rank test comparing to the hypothetical value of 150. F-G) Tactile sensitivity measurements day 1-9 (basal response day 1 is shown in D). Sensitivity was measured F) acutely 2 hrs. post LEV or VEH administration and G) prior to drug injections. Data are presented means ± SEMs and calculated as 50% withdrawal threshold (g) after square root transformation (SQRT). Mixed-effects analysis with Dunnett’s correction for multiple comparison (* = compared to positive control (WT LEV)). H-J) Dilation of the common carotid artery shown as percent of precontraction (U46619 0.1 µM) to increasing concentrations of H) Lev (0.1 µM, 1 µM, 10 µM and 30 µM), I) Carbachol (0.01 µM, 0.1 µM, 1 µM and 10 µM) and J) vehicle (0.001%, 0.01%, 0.1% and 0.3% DMSO) from WT and *eNOS^-/-^* mice. Relaxation is calculated as % of 100 precontraction. Data are presented as individual data points with mean ± SEM. Two-way ANOVA with *post hoc* Šídák analysis for multiple comparison comparing WT and *eNOS^-/-^* within each concentration. * *= P* < 0.05, ** *= P* < 0.01, *** *= P* < 0.001, **** *= P* < 0.0001.

Carotid arteries isolated from *eNOS^-/-^* mice had impaired dilatory responses to levcromakalim. These arterial segments dilated −2.8 ± 0.81%, 3.6 ± 2.0%, 13.3 ± 2.9% and 16.6 ± 4.2% in response to increasing doses (0.1 µM, 1 µM, 10 µM and 30 µM) of levcromakalim compared to −3.8 ± 0.87%, 28.8 ± 2.7%, 44.1 ± 3.1% and 59.6 ± 3.0 % dilation in arteries from WT mice, P < 0.0001 (Fig 4H). Furthermore, arteries from *eNOS^-/-^* mice had complete loss of response to acetyl cholin receptor agonist carbachol (0.01 µM, 0.1 µM, 1 µM, 10 µM) where mean dilation was −2.5 ± 0.73%, −3.8 ± 1.6%, 6.6 ± 3.0% and −3.6 ± 2.8% as compared to −1.1 ± 2.7%, 40.4 ± 6.9%, 71.2 ± 1.5% and 42.8 ± 4.5% in WT controls, P < 0.0001 (Fig 4I). This serves as a functional control for eNOS deletion [38], and suggests that the reminiscent dilation to levcromakalim is not mediated by residual eNOS activity. Vehicle exposure (0.001%, 0.01%, 0.1% and 0.3% DMSO) caused no dilation of arteries (*P =* 0.1853) (Fig 4J).

### Chemical inhibition of iNOS partially inhibited levcromakalim-induced hypersensitivity without vascular effect

As genetically knockout of eNOS did not completely block levcromakalim-induced hypersensitivity as compared to L-NAME and gene expression indicated upregulation of iNOS in dura mater post levcromakalim treatment, we tested the specific iNOS blocker *S-* methylisothiourea (SMT) in the behavioral mouse model (Fig 5A). At baseline, all treatment groups had equal 50% withdrawal thresholds: SMT+VEH (0.883 ± 0.054), SMT+LEV (0.8762 ± 0.048) and VEH+LEV (0.8758 ± 0.059) (P > 0.9830) (Fig 5B). Note that these baseline measurements are generally lower than baseline measurement in the other VF experiments which ranged from 1.065 g to 1.204 g. At the 2 h timepoint, we found no clear treatment effect (P = 0.1255) but found the negative (SMT + VEH) and positive control groups (VEH + LEV) to differ on day 9 with 50% withdrawal thresholds of 0.55 ± 0.058 and 0.24 ± 0.055 respectively (P = 0.0012). The SMT + LEV group followed the negative control group and was also significantly different from the LEV group on day 9 with 50% withdrawal thresholds of 0.54 ± 0.060 (P = 0.0020) (Fig 5C). Despite challenges with the control groups decreasing the resolution and window for observing an effect of SMT, blocking with SMT seemed to prevent development of hypersensitivity after levcromakalim as this test group followed the negative control group. Note that this is a cautious interpretation of the data. At the basal measurements, there was no overall difference between the negative (SMT + VEH) and positive control groups (VEH + LEV) (P = 0.2442) making it impossible to determine the effect of SMT on the basal hypersensitivity. On day 7, there was significant difference between the negative and positive control groups (P = 0.0034) as revealed by the posthoc analysis (Fig 5D). Lastly, motor performance assessment using rotarod showed that LEV in combination the VEH and SMT seemed to impair motor function with P values of P = 0.0156 for both groups, but SMT in combination with VEH was not significantly different. LEV was tested in all other experiments as well, without impairing motor function. Median values are 150 s for all treatment groups (Fig 5E). The few mice with poor motor performance in this experiment, did not differ from the group mean in the VF dataset. Thus, we consider this a coincidental finding and argue that is has not influenced the VF data collection.

**Figure 5:**
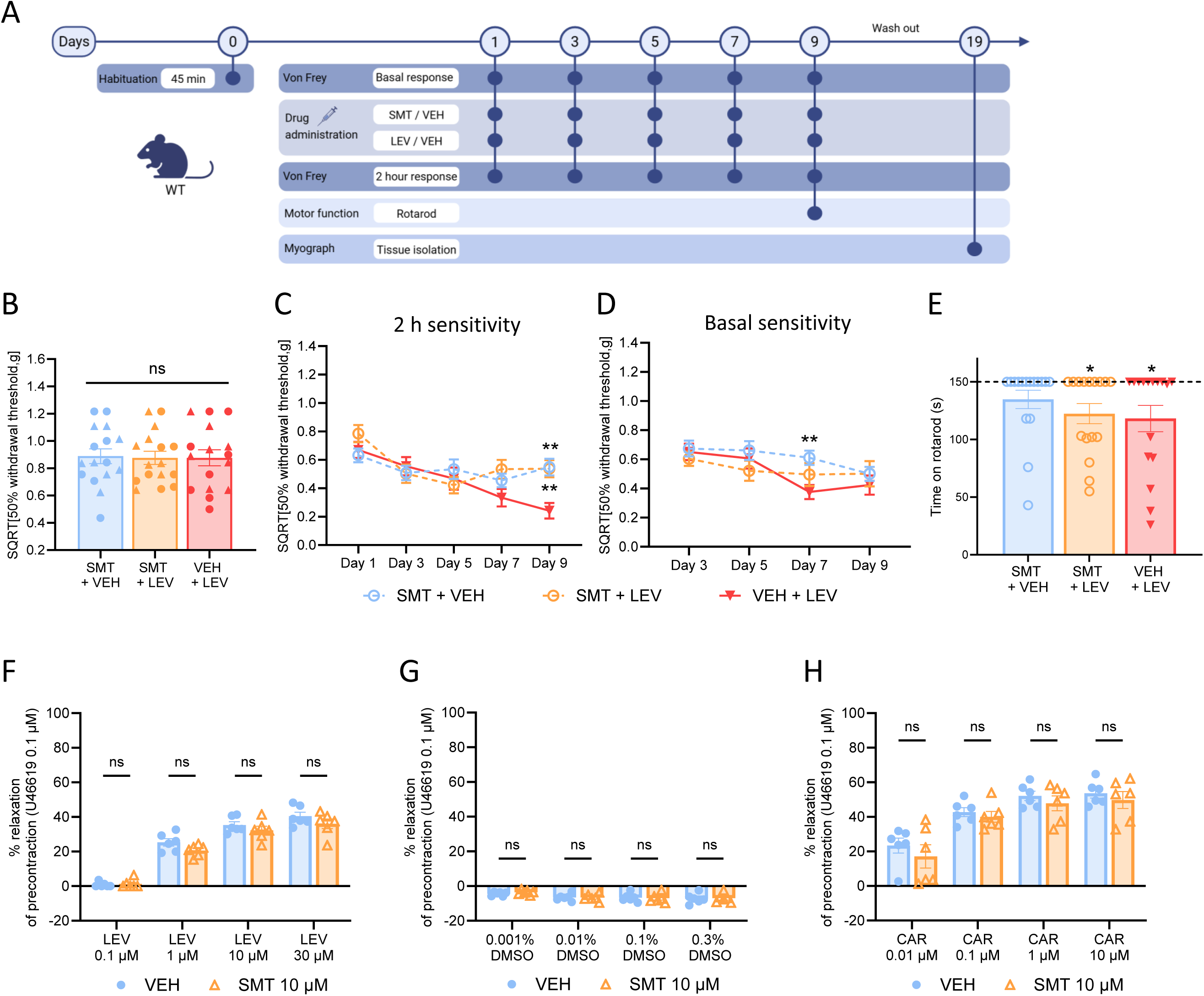
Effect of chemical inhibition of iNOS by SMT on levcromakalim-induced hypersensitivity. A) Design and timeline of the test paradigm for levcromakalim (LEV)-induced hypersensitivity and blocker S-methylisothiourea (SMT) administration. One day before drug administration, mice were placed in the test chambers for 45 min (Day 0 – Habituation). Cutaneous tactile sensitivity measurements on the plantar surface of the left hind paw were performed using von Frey (VF) filaments on every other day before (Basal response) and 2 hours (2 h response) after drug administration for a total of 9 days. Before each VF test, mice were placed in test chambers 30-45 min prior to testing. On every test day, animals received SMT (10 mg/kg i.p.) 15-20 minutes prior to LEV (1 mg/kg i.p.) administration. Following the final VF test on day 9, motor function was evaluated by rotarod performance. After a washout period of 10 days, common carotid arteries were isolated for myography experiments. The illustration is created with Biorender.com. B) Baseline tactile sensitivity values on day 1. Data are presented as individual data points with triangles representing males (▲) and circles representing females (●). One-way ANOVA with *post hoc* Tukey multiple comparison test. C-D) Tactile sensitivity measurements day 1-9 (basal response day 1 is shown in B). Sensitivity was measured C) acutely 2 hrs. post LEV or VEH administration and D) prior to drug injections. Data are presented means ± SEMs and calculated as 50% withdrawal threshold (g) after square root transformation (SQRT). Mixed-effects analysis with Dunnett’s correction for multiple comparison (* = compared to positive control (VEH+LEV)). E) Rotarod performance. Data are shown as individual data points with mean ± SEM. Wilcoxon signed-rank test comparing to the hypothetical value of 150. F-H) Dilation of the common carotid artery shown as percent of precontraction (U46619 0.1 µM) to increasing concentrations of F) Lev (0.1 µM, 1 µM, 10 µM and 30 µM), G) vehicle (0.001%, 0.01%, 0.1% and 0.3% DMSO) and H) Carbachol (0.01 µM, 0.1 µM, 1 µM and 10 µM) from WT mice treated with 10 µM SMT or vehicle. Relaxation is calculated as % of 100 precontraction. Data are presented as individual data points with mean ± SEM. Two-way ANOVA with *post hoc* Šídák analysis for multiple comparison comparing SMT and VEH within each concentration. * *= P* < 0.05, ** *= P* < 0.01, *** *= P* < 0.001, **** *= P* < 0.0001.

Isolated carotid arteries from WT mice treated with SMT 10 µM had the same dilatory responses to increasing concentrations of levcromakalim as vehicle-treated arteries (P values ranging from 0.34 to 0.99) (Fig 5F). Specifically, at concentrations of 0.1 µM, 1 µM, 10 µM and 30 µM levcromakalim, carotid artery dilation in SMT-treated arteries was 0.8 ± 0.6%, 25.3 ± 2.1%, 35.3 ± 1.9%, and 40.4 ± 2.3%, respectively, compared to 1.3 ± 1.0%, 20.6 ± 1.4%, 32.2 ± 2.4% and 36.1 ± 2.8% in vehicle-treated arteries. Vehicle exposure (0.001%, 0.01%, 0.1% and 0.3% DMSO) caused no dilation of arteries (*P* = 0.8767) (Fig 5G). To assess endothelial function, response to acetyl cholin receptor agonist carbachol (0.01 µM, 0.1 µM, 1 µM, 10 µM) was tested. There was no difference between SMT and vehicle-treated arteries with mean dilation of SMT-treated arteries being 23.4 ± 4.3%, 42.8 ± 2.6%, 52.2 ± 2.7% and 53.7 ± 2.8% as compared to 17.1 ± 6.7%, 40.1 ± 3.1%, 47.9 ± 4.3% and 49.8 ± 4.9% in vehicle-treated arteries, P = 0.3761 (Fig 5H).

### NO produced in response to levcromakalim engaged reactive oxygen/nitrogen pathways rather than the sGC pathway to induce hypersensitivity

Given the eNOS and possibly iNOS involvement in levcromakalim-induced hypersensitivity, we further explored downstream pathways of NO. The sGC inhibitor ODQ 1 mg/kg (i.p.) was effective against GTN-induced hypersensitivity [1], and thus ODQ 1 mg/kg was first tested in the levcromakalim model (Fig 6A). Observing no effect of ODQ 1 mg/kg (Fig 6B-D), we increased the dose to 10 mg/kg (i.p.) (Fig 6E-G) but found that this dose also did not inhibit levcromakalim-induced hypersensitivity, neither at the acute nor basal measures. Pre-group allocation baseline measurements were equal among all treatment groups in both cohorts (ODQ 1mg/kg; *P* = 0.9790, ODQ 10mg/kg; *P* = 0.9811) (Fig 6B and E). In both experiments ODQ+LEV-treated mice had 50% withdrawal thresholds equal to or lower than VEH+LEV-treated mice on all test days. Overall, VEH-VEH and ODQ-VEH control groups never had 50 % withdrawal thresholds below 0.9 whereas VEH+LEV and ODQ+LEV progressively dropped below this reaching a minimum of 0.42, P = 0.07-<0.0001 (days 3-9 2 h and basal). Based on this, it is unlikely that NO generated by eNOS and possibly iNOS in response to levcromakalim targets sGC to drive mechanical hypersensitivity.

**Figure 6:**
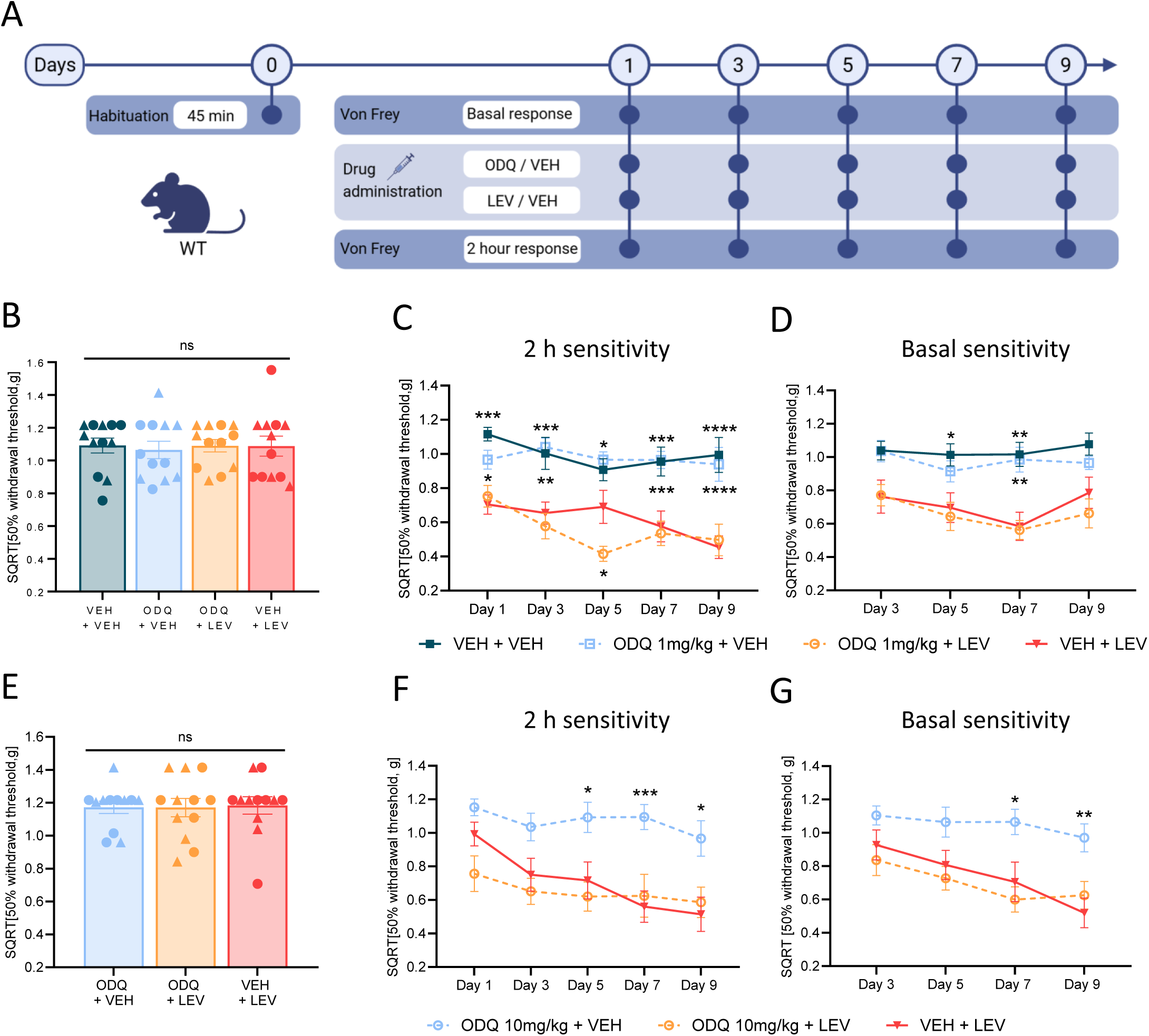
Effect of chemical inhibition of soluble guanylate cyclase by ODQ on levcromakalim-induced hypersensitivity. A) Design and timeline of the test paradigm for levcromakalim (LEV)-induced hypersensitivity and blocker 1H-[1,2,4]oxadiazolo[4,3-a]quinoxalin-1-one (ODQ) administration at two different doses. One day before drug administration, mice were placed in the test chambers for 45 min (Day 0 – Habituation). Cutaneous tactile sensitivity measurements on the plantar surface of the left hind paw were performed using von Frey (VF) filaments on every other day before (Basal response) and 2 hours (2 h response) after drug administration for a total of 9 days. Before each VF test, mice were placed in test chambers 30-45 min prior to testing. On every test day, animals received ODQ (1 or 10 mg/kg, i.p) 15-20 minutes prior to LEV (1 mg/kg, i.p.) administration. The illustration is created with Biorender.com. B) Baseline tactile sensitivity values on day 1 prior to ODQ low dose experiment. Data are presented as individual data points with triangles representing males (▲) and circles representing females (●). One-way ANOVA with *post hoc* Tukey multiple comparison test. C-D) Tactile sensitivity measurements day 1-9 for ODQ low dose (basal response day 1 is shown in B). Sensitivity was measured C) acutely 2 hrs. post LEV or VEH administration and D) prior to drug injections. Data are presented means ± SEMs and calculated as 50% withdrawal threshold (g) after square root transformation (SQRT). Mixed-effects analysis with Dunnett’s correction for multiple comparison (* = compared to positive control (VEH+LEV)). E) Baseline tactile sensitivity values on day 1 prior to ODQ high dose experiment. Data are presented as individual data points with triangles representing males (▲) and circles representing females (●). One-way ANOVA with *post hoc* Tukey multiple comparison test. F-G) Tactile sensitivity measurements day 1-9 for ODQ high dose (basal response day 1 is shown in E). Sensitivity was measured F) acutely 2 hrs. post LEV or VEH administration and G) prior to drug injections. Data are presented means ± SEMs and calculated as 50% withdrawal threshold (g) after square root transformation (SQRT). Mixed-effects analysis with Dunnett’s correction for multiple comparison (* = compared to positive control (VEH+LEV)). * *= P* < 0.05, ** *= P* < 0.01, *** *= P* < 0.001, **** *= P* < 0.0001.

Therefore, we investigated possible involvement of PN generation and found that the PN decomposition catalyst FeTPPS, was partially effective against levcromakalim-induced hypersensitivity (Fig 7A). At baseline, all treatment groups had equal 50% withdrawal thresholds: FeTPPS+VEH (1.12 ± 0.046), FeTPPS+LEV (1.11 ± 0.086) and VEH+LEV (1.17 ± 0.048) (P = 0.7555) (Fig 7B). At the 2 h time point, levcromakalim induced hypersensitivity in the positive control group (VEH+LEV) from day 3-9 (P = <0.0001-0.001). FeTPPS partially blocked levcromakalim-induced hypersensitivity with mean differences in SQRT 50% withdrawal thresholds between the FeTPPS+LEV group and the VEH+LEV group was 0.17-0.31g (days 1-9), which were significant day 5, 7, and 9, P = 0.008-0.02 (Fig 7C). FeTPPS did not influence the basal sensitivity measure, likely reflecting drug washout (Fig 7D). Lastly, FeTPPS did not affect locomotor activity as examined by rotarod performance (*P* values > 0.1250, MIDA; *P* = 0.0039) (Fig 7E). These results show that hypersensitivity induced by levcromakalim is to some degree mediated via PN generation.

**Figure 7:**
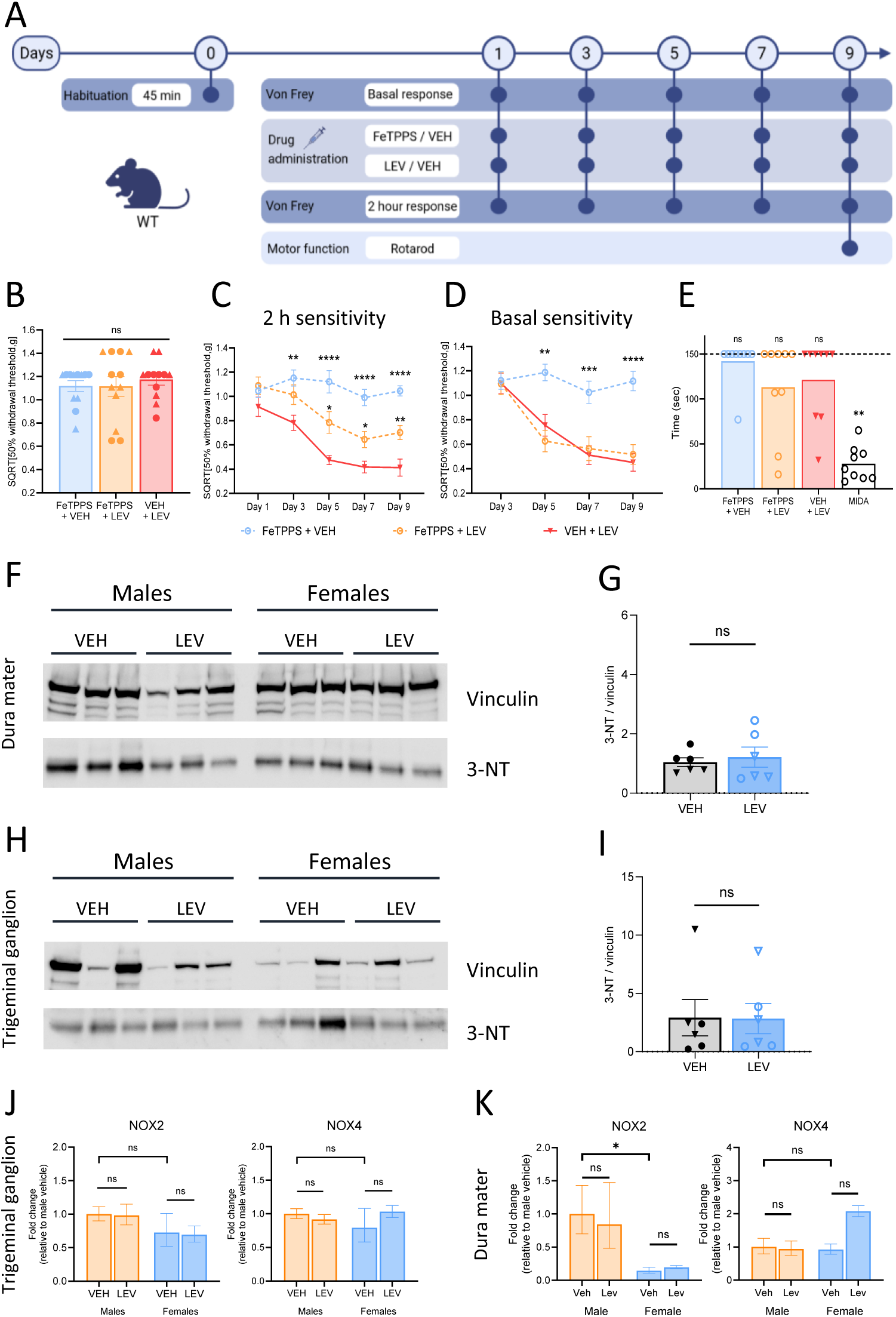
Effect of peroxynitrite decomposition catalyst FeTPPS on levcromakalim-induced hypersensitivity. A) Design and timeline of the test paradigm for levcromakalim (LEV)-induced hypersensitivity and blocker Fe(III)5,10,15,20-tetrakis(4-sulfonatophenyl) porphyrinato chloride (FeTPPS) administration. One day before drug administration, mice were placed in the test chambers for 45 min (Day 0 – Habituation). Cutaneous tactile sensitivity measurements on the plantar surface of the left hind paw were performed using von Frey (VF) filaments on every other day before (Basal response) and 2 hours (2 h response) after drug administration for a total of 9 days. Before each VF test, mice were placed in test chambers 30-45 min prior to testing. On every test day, animals received FeTPPS (30 mg/kg i.p.) 15-20 minutes prior to LEV (1 mg/kg, i.p.) administration. Following the last VF test on day 9, motor function was evaluated by rotarod performance with midazolam (MIDA, 2 mg/kg i.p.) as a positive control. The illustration is created with Biorender.com. B) Baseline tactile sensitivity values on day 1. Data are presented as individual data points with triangles representing males (▲) and circles representing females (●). One-way ANOVA with *post hoc* Tukey multiple comparison test. C-D) Tactile sensitivity measurements day 1-9 (basal response day 1 is shown in B). Sensitivity was measured C) acutely 2 hrs. post LEV or VEH administration and D) prior to drug injections. Data are presented means ± SEMs and calculated as 50% withdrawal threshold (g) after square root transformation (SQRT). Mixed-effects analysis with Dunnett’s correction for multiple comparison (* = compared to positive control (VEH+LEV)). E) Rotarod performance. Data are shown as individual data points with mean ± SEM. Wilcoxon signed-rank test comparing to the hypothetical value of 150. F, H) Western blot analysis of 3-NT level in F) dura mater and H) trigeminal ganglia (TG) from mice treated with LEV (1 mg/kg i.p.) or vehicle (2 % DMSO in saline i.p.) every other day for 9 days (total of 5 administrations) as in VF hypersensitivity experiments. Tissue is from an independent cohort of mice. Protein levels were normalized to house-keeping gene vinculin. G, I) Quantification of western blots shown in (F) and (H). Quantification of 3-NT level in from G) dura mater and I) TG from mice treated with LEV (1 mg/kg i.p.) or vehicle (2 % DMSO in saline i.p.) every other day for 9 days (total of 5 administrations). Data are presented as mean ± SEM. Unpaired t-tests. J-M) Gene expression of *Nox2* and *Nox4* in J) TG and K) dura mater from mice treated with LEV (1 mg/kg i.p.) or vehicle (2 % DMSO in saline i.p.). Gene expression was normalized to *B-actin*. Fold change is shown relative to male vehicle and data are presented as mean ± SEM. One-way ANOVA with Šídák’s correction for multiple comparison. * *= P* < 0.05, ** *= P* < 0.01, *** *= P* < 0.001, **** *= P* < 0.0001.

To further investigate PN activation, we measured 3-nitrotyrosine (3-NT), a well-established marker of PN activity, as it reflects protein nitration, which can alter function and contribute to neuronal excitability [32,54,55]. Given that accelerating PN breakdown partially blocks levcromakalim-induced hypersensitivity, we hypothesized that repeated levcromakalim administration would elevate 3-NT expression. We measured 3-NT expression in the TG and dura mater following a full paradigm of repeated administration of levcromakalim. However, western blot analysis did not reveal a significant increase in 3-NT expression in neither the dura mater (Fig 7F and G) nor TG (Fig 7H and I), with no significant differences between VEH- and LEV-treated groups (Dura mater; P = 0.652, TG; P = 0.968).

### Explorative gene expression analysis did not reveal changes in NOX expression following levcromakalim administration

PN is generated through the reaction of superoxide (O₂⁻) and nitric oxide (NO), with superoxide primarily produced by NADPH oxidases (NOX enzymes). Given that eNOS- and possibly iNOS-derived NO and PN formation are implicated in levcromakalim-induced hypersensitivity, we conducted an exploratory analysis of NOX mRNA expression following repeated levcromakalim administration. Specifically, we assessed the expression of NOX2 and NOX4 in TG and dura mater (Fig 7J). In TG, levcromakalim treatment did not significantly alter NOX2 (Males; *P* > 0.9999, Females; *P* = 0.9987) or NOX4 (Males; *P* = 0.9830, Females; *P* = 0.6970) (Fig 7J) expression, with no differences observed between males and females (NOX2; *P* = 0.7074, NOX4; *P* = 0.7684. In dura mater, neither NOX2 (Males; *P* > 0.9999, Females; *P* = 0.9321) nor NOX4 (Males; *P* > 0.5283, Females; *P* = 0.2632) mRNA levels were altered (Fig 7K). Although, there was a tendency of an increase of NOX4 in females. Furthermore, we observed a significantly lower expression level of NOX2 in females compared to males (P = 0.0073). These findings suggest potential sex-dependent differences in NOX expression in the dura mater, which may have implications for oxidative stress regulation. However, given the exploratory nature of this analysis, further investigation is required ideally on protein level. We also intended to measure NOX1 expression in TG and dura mater but found expression to be below detection limit.

## Discussion

The mechanism of headache and migraine induction following direct and indirect opening of K_ATP_ channels is largely unexplored but is becoming a topic of increased interest [16,20,21,48,56]. Here, we show that engagement and upregulation of NOS enzymes in the dura mater is a key process giving rise to *de novo* NO production and PN generation. Our findings *in vivo* and on isolated carotid arteries indicate a strong eNOS dependency both on levcromakalim-induced tactile hypersensitivity and vasodilation. The contribution from nNOS was minor, whereas cautious interpretation also called for a relevance of iNOS on behavioral but not vascular responses. Furthermore, we found that the classical NO-receptor sGC was not involved in the levcromakalim-induced hypersensitivity response, whereas PN-generation drove part of the phenotype. We suggest that repeated levcromakalim induces a state of both coupled and uncoupled eNOS activity. The former causing more NO to be produced and the latter stimulating oxidative stress and the formation of H_2_O_2_ and PN. The NOS induction and link to nitrosative stress can possibly explain why migraine attacks “build up” and peak hours after drug-administration and is maintained for a while although most migraine triggers and NO have short half-lifes [5,6,39,66].

### NOS across migraine models and spontaneous attacks

This is the first study to show NOS involvement in the behavioral response to levcromakalim. Within vascular physiology, NOS dependency of K_ATP_ and hyperpolarization-induced dilation is an established concept where NO produced by eNOS in endothelial cells acts on sGC in vascular smooth muscle to drive relaxation [40,46]. Furthermore, NOS involvement is described in multiple other preclinical migraine models [29,52]. In rats, CGRP as well as neurogenically induced dilation of meningeal arteries was inhibited by NOS inhibitors, GCRP-induced dilation exclusively by the eNOS inhibitors N5-(Iminoethyl)-L-ornithine and diphenyleneiodonium chloride and neurogenic dilation only by the nNOS inhibitor SMTC [3]. In rat aorta, NOS inhibitors have also been found to attenuate CGRP-induced endothelium-dependent relaxation of smooth muscle by restricting the eNOS activity in the endothelium [33]. Yet, lack of selectivity of inhibitors calls for careful interpretation. A recent study found no efficacy of L-NAME on CGRP-induced tactile hypersensitivity [2]. We found a slight effect of SMTC on levcromakalim-induced hypersensitivity, but this was not seen in *nNOS^-/-^* mice suggesting that the SMTC effect may have been contributable to minor eNOS inhibition at the selected dose. Also, nNOS did not contribute to levcromakalim-induced dilation of isolated arteries. However, in GTN models nNOS contribution to pain readouts cannot be ruled out. The inhibitor 7-nitroindazole was effective in reducing TNC c-fos levels after GTN administration in rats [65], and SMTC partly inhibited tactile hypersensitivity induced by GTN in mice but could not be confirmed in *nNOS^-/-^* mice. As we experienced challenges with the controls in our iNOS experiment, it is with caution, that we conclude that iNOS may be involved in levcromakalim-induced hypersensitivity. Yet, our results are supported by the enhanced iNOS gene expression observed in dura mater and align with other studies suggesting a relevance of iNOS in headache genesis. In mice, SMT in the same dose as applied here – prevented GTN-induced hypersensitivity [29]. Similarly, in rats, infusion of GTN induced iNOS mRNA expression in meningeal macrophages [59] and c-FOS expression was lowered by non-selective NOS inhibition [57].

Thus, in preclinical experiments NOS activity is a common denominator for at least migraine triggers GTN, CGRP, and levcromakalim suggesting that endogenous NO production is a driver of headache genesis in these models. The gene expression data from this study further support the involvement of NOS isoforms in the dura mater following administration of levcromakalim. Upregulation of both eNOS and iNOS was observed exclusively in the dura mater, with no significant changes in the TG or cerebral blood vessels. eNOS upregulation in dura mater is likely localized to the endothelial cells of the dural vasculature, while iNOS may be expressed in resident immune cells such as macrophages as previously described [59]. As the mRNA analysis was performed on whole dura mater tissue and not at the single-cell level, the exact cellular sources remain speculative. Interestingly, the slight upregulation of nNOS in the dura mater did not translate into detectable changes in the TG, highlighting the need to clarify the spatial distribution and possible functional role of nNOS in the dura mater. Overall, these data enforce the notion that the dura mater is a key site of action for migraine triggers and plays a central role in migraine pathophysiology [20,48].

Clinically, the importance of NOS enzymes in headache pathophysiology was demonstrated by efficacy of non-selective NOS inhibition in acute migraine attacks and in chronic tension-type headache [10,45]. In a large cohort, it was recently found that interictal L-arginine levels were lower in the cerebrospinal fluid (CSF) of people with migraine without aura as well as migraine with aura compared to controls [50]. L-arginine is the substrate for endogenous NO production by NOS suggesting altered NO homeostasis in people with migraine. It has not yet been possible to develop selective or non-selective NOS inhibitors for migraine. Phase II clinical trials of a selective iNOS inhibitor were unsuccessful both for acute and preventive treatment of migraine [35] suggesting that iNOS inhibition alone is insufficient. A clinical trial exploring a combined nNOS inhibitor and 5HT-1B/1D agonist showed mixed results [36]. eNOS inhibition has not been assessed clinically and is not a sensible drug target due to its fundamental role in healthy cardiovascular function [13].

### Nitrosative stress

Our findings indicate that endogenous NO produced by NOS in the levcromakalim mouse model, subsequently acts via PN generation and not its classical receptor sGC. The PN decomposition catalyst FeTPPS partially inhibited the effect of levcromakalim in the behavioral model whereas sGC blocker ODQ was ineffective. PN is a short lived, toxic anion generated by reaction of NO and superoxide (O_2_^-^) that may further react and form other reactive species [63,64]. As for NOS activity, PN generation is seen across mechanistically distinct preclinical models. In 1999, decreased levels of superoxide were shown in rats following GTN infusion and PN generation were suggested as the product [58]. Three recent migraine relevant studies showed efficacy of FeTPPS on stress + SNP-induced behavioral and biochemical readouts [44], GTN-induced tactile hypersensitivity [29], and CGRP-induced neuronal and behavioral changes [2], respectively. PN activity can be measured by oxidative stress biomarker 3-nitrotyrosine [12]. In our TG and dura mater samples from levcromakalim-treated mice we did not find elevated expression of 3-NT as seen in the restraint stress + SNP mouse model [44]. Lack of 3-NT elevation in TG and dura mater suggests that, although PN generation appears to be involved in the levcromakalim model, its presence may be temporally or spatially different than our sampling or simply below detection level.

Likewise, Marone *et al.* showed that generation of reactive oxygen and carbonyl species generated via TRPA1 and NOX in the TG drives GTN induced hypersensitivity[47] and a novel rat study suggests that Romo1 (reactive oxygen species modulator 1) is an important mediator of central sensitization in response to recurrent GTN injections [71]. We did not see upregulation of NOX mRNA in neither dura mater nor TG in response to levcromakalim. This does not mean that NOX involvement can be ruled out, only that the expression pattern is not regulated. However, activity of the enzyme is likely regulated and should be assessed by protein analysis of NOX inhibition in future studies. But we did see possible sex differences in expression patterns of NOX2 in dura mater which may be of relevance to sex specific differences in migraine pathophysiology.

Clinically, evidence of higher nitrosative and oxidative stress in people with migraine remains inconclusive but points towards possible lower levels of superoxide dismutase (SOD), higher levels of NO; nitrite/nitrate, elevated thiobarbituric acid reactive substances (TBARS) which is a marker of lipid peroxidation [49,51], and other markers. Multiple studies report reduced headache frequency and antioxidant effect of dietary supplements such as selenium [11], magnesium [23], riboflavin [70]commonly used as supplements for management of migraine [31]. However, solid evidence from large controlled clinical trials is lacking.

### Migraine loops - Interrelated pathways of headache inducing drugs

Migraine may be induced in people with migraine diagnosis by a series of trigger substances some of which have already been mentioned. These triggers are all potent vasodilators supposedly causing smooth muscle hyperpolarization via G-protein coupled receptor (GPCR)-mediated or direct cAMP and/or cGMP accumulation ultimately opening K_ATP_ channels and/or large conductance calcium-activated potassium (BK) channels [4]. The simplicity of this up-down signaling axis is challenged by findings showing that one vasodilator may induce production/release of other vasodilators as is also the case here. In other studies, it was shown that increased levels of endogenous NO were measured in rats during GTN infusion [58], NO is linked to the release of CGRP [25], dilation of meningeal arteries induced by CGRP is inhibited by NOS inhibitors [3], and nociceptive responses in mice to K_ATP_ channel opener levcromakalim is CGRP dependent [21]. Thus, several of the vasodilating migraine triggering substances may have interrelated pathways and loops potentiating the effect of one-another. We have previously described a K_ATP_-CGRP pathway [21] which may likely be linked to the NO loop described here, as NO and CGRP are closely related. Adding to the interrelated mechanisms, the TRPA1 ion channel which mediates CGRP release and GTN-induced hypersensitivity [22,47] is also sensitive to mediators of oxidative stress [68]. But as mentioned, we have previously described, that the mouse model of levcromakalim-induced migraine is not dependent on TRPA1 [21] stressing that the exact same mechanisms are not employed for GTN and levcromakalim-induced migraine and that we still do not fully understand the interrelated signaling pathways.

### Strengths and limitations

An important point of discussion is the validity of the repeated levcromakalim provocation model for human migraine. In human provocation experiments, only a single administration of a trigger is sufficient to precipitate headache and/or migraine attacks. In WT mice, the majority needs more than a single injection for establishment of robust tactile hypersensitivity making the mouse and human experimental models somewhat different. On the other hand, validity is strongly supported by the effect of highly specific migraine treatments olcegepant and sumatriptan and we see the applied mouse model as a useful mechanistic tool to study relevant signaling pathways *in vivo*. The major strength of our study is the comprehensive *in vivo* behavioral evidence supported by *ex vivo* vascular and gene expression data. All data and analyses are presented with focus on transparent reporting. A weakness of the study is the heterogeneity in levcromakalim-induced hypersensitivity responses between experimental cohorts and experimenters. This hampered our interpretation on iNOS involvement in the levcromakalim model. Unfortunately, repetition of the experiment was not possible. Minor weaknesses include the lack of chemical inhibitors specific for eNOS and nNOS demanding the use of global knockout mice. For the nNOS strain, knockouts and controls were not bred in parallel but purchased from different vendors. In our interpretation, this did not affect the overall conclusion of the study. Furthermore, it was not possible to conduct more experiments to make stronger interpretations on specific cell type involvement and NOX activity.

### Conclusion and future perspectives

Our study found that tactile hypersensitivity induced by K_ATP_ channel opener levcromakalim in mice is driven by NOS activity and peroxynitrite generation in the dura mater.

The study highlights the complex interplay between K_ATP_ channel activation, NOS isoform regulation, and nitrosative stress in migraine pathophysiology. Future research should aim to delineate the precise signaling events following endothelial and smooth muscle hyperpolarization, particularly during the lag phase preceding migraine onset. Moreover, the spatial and temporal dynamics of PN activity and its interaction with other oxidative stress pathways as well as the differential roles of NOS isoforms in distinct cellular compartments of the dura mater warrant deeper investigation. Finally, given the interrelated nature of migraine triggers such as CGRP, NO, and K_ATP_ channel openers, future studies should explore how these pathways converge or diverge. NOS may not be a relevant drug target, but the wide range of other targets such as PN, NOX, or SOD may be feasible either as monotherapy but more likely in addition to other treatment modalities with suboptimal effect.

## Acknowledgements

This section is not mandatory.

