## Supplementary material for "*De novo* nitric oxide synthesis drives tactile hypersensitivity induced by ATP-sensitive potassium channel opening in mice: Relevance to migraine and other headache disorders"

**Genotyping**

*eNOS^-/-^* and corresponding WT mice bred in-house were continuously monitored through genotyping. WT animals received from Taconic and *nNOS^-/-^* mice from JAX were not genotyped.

*DNA extraction*

Animals were ear marked upon weaning, and DNA was extracted from those samples. To each tube, 100 µL Alkaline Lysis Buffer (25 mM NaOH + 0.25 mM EDTA in MiliQ H_2_O) was added and then spun down (11,0 g for 30 sec, Eppendorf MiniSpin plus centrifuge, Eppendorf, Germany) to ensure tissue submersion. Samples were boiled at 95 °C for 30 min (Stuart SBH200D Block Heater, Cole-Parmer, IL, USA) before adding 100 µL Neutralization Buffer (40 mM Tris HCl in MiliQ H_2_O) and vortexed. Supernatant now contained DNA and was either frozen for later or put on ice for immediate use.

*PCR*

Samples were then prepared to polymerase chain reaction (PCR). Master mix was prepared by mixing RNase free H_2_O, RedTaq ReadyMix (Sigma Aldrich, MO, USA), and the three primers used in the JAX genotype protocol (<https://www.jax.org/Protocol?stockNumber=002684&protocolID=23416>), produced by TAG Copenhagen (Denmark). Protocol was optimized in-house and therefore deviate from the JAX protocol. Master mix reagents and PCR program is summarized in Tables S1 and S2.

Table S1. Overview of master mix components for eNOS genotyping.

| Master mix | |
| --- | --- |
| Reagents | Volume pr. reaction |
| Primer 1 (10 µM) | 0.67 µL |
| Primer 2 (10 µM) | 0.67 µL |
| Primer 3 (10 µM) | 0.67 µL |
| Redtaq ReadyMix | 10 µL |
| RNase free H_2_O | 5.99 µL |
| Total | 18 µL (+2 µL DNA) |

Table S2. PCR-program for eNOS genotyping.

| Step | Temperature | Time | Cycles |
| --- | --- | --- | --- |
| Step 1: Initial Denaturation | 94 °C | 2 min | 1 |
| Step 2: |  |  |  |
| Denaturation | 94 °C | 30 s |  |
| Annealing | 59 °C | 45 s | 35 |
| Elongation | 72 °C | 30 s |  |
| Step 3: Final Elongation | 72 °C | 10 min |  |
| Step 4: Storage | 4 °C | ∞ |  |

*Gel electrophoresis*

A 1% agarose in 1X TAE buffer gel was prepared with GelGreen Nucleic Acid Gel Stain (Biotium, CA, USA). PCR products were then run in 1X TAE buffer for 55-75 min at 50-80V and viewed on LAS-4000 Imager (Fujifilm, Japan).

**RNA extraction for qRT-PCR**

Extracted tissues were kept at -80°C until needed. Working in a fume hood, 2 ml FastPrep tubes were filled to about half with ceramic beads and 1 ml Trizol was added before added tissue samples. Using a Fastprep (FastPrep-24 5G, MP Biomedicals, USA), tissues was homogenized at settings 6 m/s, 3x20 s with 20 s breaks in-between. Samples were then left on ice for around 10 minutes to settle before being transferred to RNase free tubes taking care not to transfer foam. To each tube, 0.2 ml chloroform was added, mixed, and centrifuged at 4°C for 20 min (12,000 x g, Eppendorf centrifuge 5417R, Eppendorf, Germany). The upper, aqueous phase was carefully extracted and transferred to a new tube. An equal volume isopropanol was added, mixed, and left on bench for around 10 min. Samples were then spun for 30 min (4°C, 12,000 x g) before supernatant was carefully removed and discarded. Last traces of supernatant were removed by centrifuging a one more time (4°C, 3 min, 12,000 x g). RNA pellet was then cleaned by adding 1 ml of ice-cold 75% ethanol, inverting tubes until pellet is free-floating and then spun for 15 min (4°C, 12,000 x g). Supernatant was removed, samples were spun again (4°C, 12,000 x g), and supernatant traces were removed carefully without disturbing RNA pellet. Tubes were left for approximately 20 min on bench with open lids to allow evaporation of ethanol traces. RNA pellet was then eluted in an appropriate volume (RNase free H_2_O, 20-50 µL), vortexed and kept at 80°C. RNase Zap was repeatedly used throughout protocol to clean hood, pipettes, racks, gloves etc.

**Protein isolation from trizol samples**

After RNA had been extracted, 100µL chloroform was added to the samples. The trizol supernatant was transferred to a new tube following 15 min of centrifugation (12,000 x g, Eppendorf centrifuge 5417R, Eppendorf, Germany). Any remaining aqueous phase was removed and 240µL of 100% ethanol was added to each sample. Following 2-3 min of incubation, samples were centrifuged for 5 min at 4°C (2,000 x g, Eppendorf centrifuge 5417R, Eppendorf, Germany) and supernatant was transferred to a new tube. 1.5 ml acetone and mix by inversion. Following 10 min of incubation, samples were centrifuged for 10 min at 4 °C (12,000 x g, Eppendorf centrifuge 5417R, Eppendorf, Germany). Samples were washed 3 times in protein wash 1 (300 mM guanidine hydrochloride in 95% ethanol + 2.5% glycerol (V:V)) 1ml protein wash 1 is added to the sample and incubated for 10 min following by 5 min centrifugation (8,000 x g, Eppendorf centrifuge 5417C, Eppendorf, Germany). The supernatant is removed, and the washing step is repeated twice. The pellet is dispersed in protein wash 2 (2.5% glycerol (V:V) in 95% ethanol) and incubated for 10 min following by 5 min centrifugation at 4°C (8,000 x g, Eppendorf centrifuge 5417R, Eppendorf, Germany). The supernatant is removed, and the pellet left to dry for 7-10 min. The pellet is resuspended in protein solubilization buffer (50mM tris base, 150mM NaCl, 2% SDS (W:V), pH: 7.5) (20µL for dura mater and 50µL for TG) and incubated at 50°C on a heating block with rotation for 2 h. Lastly, the sample is centrifuged at 4°C for 15 min (14,000 x g, Eppendorf centrifuge 5417R, Eppendorf, Germany) and the supernatant is transferred to a new tube before performing western blotting.
